# Simultaneous inference of parental admixture proportions and admixture times from unphased local ancestry calls

**DOI:** 10.1101/2022.01.05.475139

**Authors:** Siddharth Avadhanam, Amy L. Williams

**Affiliations:** Department of Computational Biology, Cornell University, Ithaca, NY 14853, USA

## Abstract

Population genetic analyses of local ancestry tracts routinely assume that the ancestral admixture process is identical for both parents of an individual, an assumption that may be invalid when considering recent admixture. Here we present Parental Admixture Proportion Inference (PAPI), a Bayesian tool for inferring the admixture proportions and admixture times for each parent of a single admixed individual. PAPI analyzes unphased local ancestry tracts and has two components models: a binomial model that exploits the informativeness of homozygous ancestry regions to infer parental admixture proportions, and a hidden Markov model (HMM) that infers admixture times from tract lengths. Crucially, the HMM employs an approximation to the pedigree crossover dynamics that accounts for unobserved within-ancestry recombination, enabling inference of parental admixture times. We compared the accuracy of PAPI’s admixture proportion estimates with those of ANCESTOR in simulated admixed individuals and found that PAPI outperforms ANCESTOR by an average of 46% in a representative set of simulation scenarios, with PAPI’s estimates deviating from the ground truth by 0.047 on average. Moreover, PAPI’s admixture time estimates were strongly correlated with the ground truth in these simulations (*R* = 0.76), but have an average downward bias of 1.01 generations that is partly attributable to inaccuracies in local ancestry inference. As an illustration of its utility, we ran PAPI on real African Americans from the PAGE study (*N* = 5, 786) and found strong evidence of assortative mating by ancestry proportion: couples’ ancestry proportions are closer to each other than expected by chance (*P <* 10^−6^), and are highly correlated (*R* = 0.87). We anticipate that PAPI will be useful in studying the population dynamics of admixture and will also be of interest to individuals seeking to learn about their personal genealogies.

## Introduction

Widespread consumer interest in one’s personal genetic ancestry and genealogy has fueled the rise of direct-to-consumer genetic testing companies^1,2^, enabling high resolution studies of population structure, admixture, and migration patterns^3–5^. One of the most popular features these companies provide is ancestry estimation: the geographic locations of the populations from which customers inherited their DNA. The techniques for such estimation, first developed by academic researchers^6–8^, have applications both to demographic inference^9–11^ as well as to disease association mapping^12,13^

Inferring an individual’s genome-wide and locus-specific ancestry—i.e., local ancestry^7^—reveals not only that person’s genetic heritage, but also information about the genetic ancestry of his or her parents^14,15^. Indeed, each parent transmits half of his or her genome to a child, so in principle, one can recover half the DNA of each parent by analyzing data from a single individual. In practice however, phasing errors and uncertain interchromosomal phase pose substantial challenges to reconstructing the parents’ genomes, although methods that analyze multiple siblings and/or use population allele frequencies can be effective^16–18^.

Many past studies of admixed individuals have focused on population-level ancestry questions, including migration times^5,9,19^, yet analyzing data from couples has the power to provide refined information about demographic patterns. For example, it makes possible studies of assortative mating by ancestry^20–22^, could help identify subgroups with different admixture histories and may detect outlier individuals with patterns distinct from the bulk of the population.

The features derivable from local ancestry signals include population ancestry proportions and estimated times since admixture based on tract lengths^9,23^. Individual-level ancestry proportions are generally reliable to infer, depending on the ancestral populations^7,24,25^, yet computing admixture times for an individual (or his/her parents) from only one genome has not received careful attention and may be difficult. In particular, high variance in tract lengths^9^ and the finite genome size can lead to noisy admixture times estimates^26^. Even so, noisy estimates may still be useful when analyzed in aggregate across many samples.

Although phase accuracy remains an issue in the context of local ancestry inference^13^, homozygous ancestry regions provide unambiguous information about the ancestry of the two parents at that locus. As such, even when analyzing unphased local ancestry tracts^7,25^, one can leverage these homozygous regions (as well as the lack thereof) to estimate the parental ancestry fractions. Analyzing unphased local ancestry tracts also helps prevent biases due to phase inaccuracy, both in the estimating parental ancestry proportions and time since admixture (see Methods).

We developed Parental Ancestry Proportion Inference (PAPI), a tool that provides estimates of both the admixture proportions and times since admixture of the two parents of individual two-way admixed samples. PAPI uses both a binomial model and a hidden Markov model (HMM) to infer these parental values, and, by default, combines these two models into a composite likelihood. We validated PAPI using simulated and real African Americans and find that it provides reliable parent ancestry proportion estimates. PAPI’s time since admixture estimates are also well correlated with the ground truth, and it is well powered to determine when the two parents have different times. We applied PAPI to real African Americans from the PAGE study^27^, finding strong evidence for ancestry-based assortative mating (Results).

## Methods

PAPI is designed to infer the ancestry proportions (*p*_*A*_, *p*_*B*_) and the average number of generations to unadmixed ancestors (*t*_*A*_, *t*_*B*_) for the unsexed parents *A* and *B* of a single admixed individual (Figure 1). It takes unphased diploid local ancestry tracts for focal admixed individuals as input and works on two-way admixed samples, but the approach can be extended to handle multi-way admixture. Throughout this paper, we label the two populations being analyzed arbitrarily as 0 and 1.

**Figure 1:**
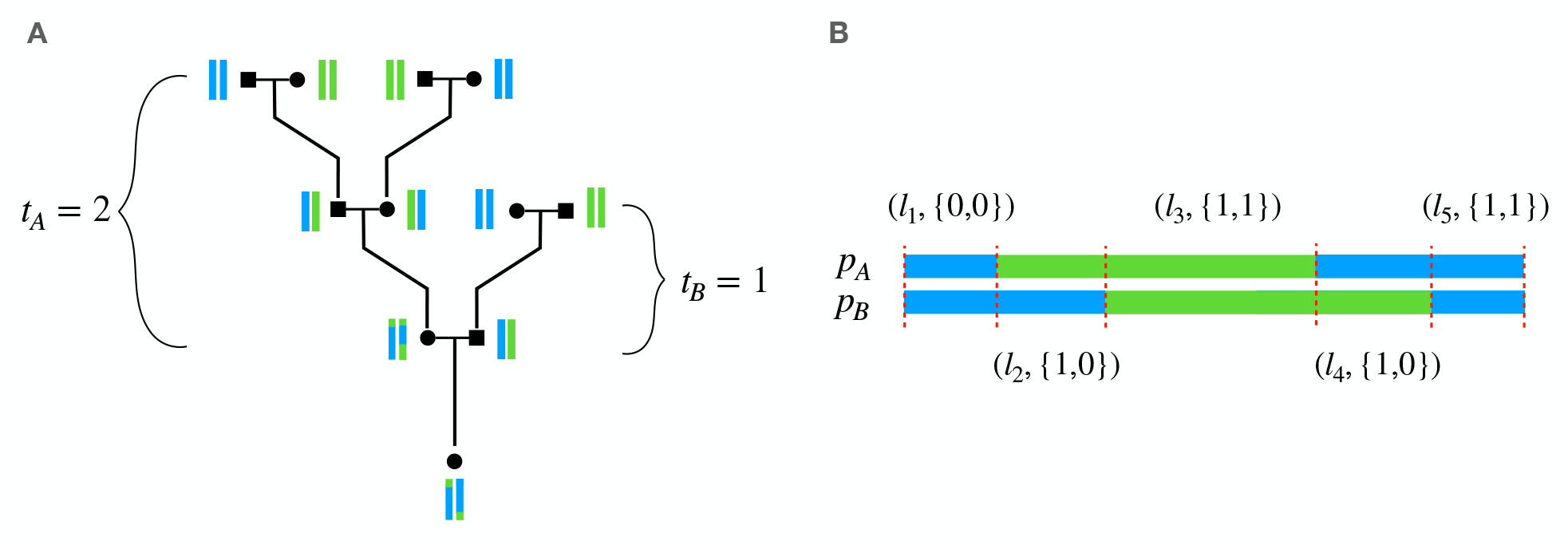
Example pedigree and local ancestry data. (A) Pedigree with vertical bars next to each individual representing two of their chromosomes, each colored to represent local ancestry from two populations, blue and green. *t*_*A*_ and *t*_*B*_ are the number of generations to the unadmixed ancestors for parents A and B, respectively; this is the generation in which at least one couple contains individuals of different ancestries. (B) PAPI takes in unphased local ancestry tracts denoted by ***x***_*i*_ = (*l*_*i*_, *a*_*i*_). The image depicts a particular phasing of the data with labels that correspond to the observed data. As depicted, PAPI’s binomial model uses a parameterization that assigns *p*_*A*_ and *p*_*B*_ to the transmitted parental haplotypes *A* and *B* respectively.

The advantage of using unphased local ancestry calls is that they are immune to switch errors that affect the phased input to (and therefore output of) haplotype-based local ancestry methods. PAPI makes inferences of times since admixture (*t*_*A*_, *t*_*B*_) based on local ancestry tract lengths, and switch errors can create artificially short or long ancestry tracts that would bias this approach if it relied on phased input. Additionally, switch errors in phased input can put an ancestry tract on an incorrect haplotype, which would lead to incorrect (*p*_*A*_, *p*_*B*_) estimates.

PAPI employs a composite likelihood built from two component models: a binomial model that infers (*p*_*A*_, *p*_*B*_) and a hidden Markov model (HMM) that jointly infers all parameters ***θ*** = (*p*_*A*_, *p*_*B*_, *t*_*A*_, *t*_*B*_). The binomial model exploits the information in regions that are homozygous for ancestry and uses genome-wide diploid ancestry proportions to infer (*p*_*A*_, *p*_*B*_). In turn, the HMM estimates ***θ*** by integrating over all possible phasings of the input local ancestry tracts using the forward algorithm. Our empirical findings indicate that forming a composite likelihood from these two models provides more accurate estimates of than those of each component alone (Results).

A key challenge in inferring (*t*_*A*_, *t*_*B*_) is that a subset of the crossovers the ancestors transmitted typically occur between haplotypes of the same ancestry, and, as a result, are not observable in local ancestry calls. The HMM accounts for this by approximating the rate of these hidden crossovers from the parental ancestry proportions (*p*_*A*_, *p*_*B*_) as described below. This approximation relies on a population genetic model of ancestry tract length distributions following a pulse migration^9^, which serves here as a simplified model of the complex dynamics of the meioses in the focal individual’s pedigree.

### A binomial model to infer parent ancestry proportions

The binomial model for estimating (*p*_*A*_, *p*_*B*_) operates on genome-wide diploid ancestry fractions. As a motivating example, consider a locus in the focal individual that is homozygous for ancestry. This locus carries unambiguous information about both parents’ ancestries—it is certain that each parent carries at least one copy of the given ancestry in that region. Furthermore, low proportions of homozygous ancestry regions indicate that the parents likely differ in their ancestry at most loci. For example, a child that is heterozygous for ancestry at every position likely has two parents with very high levels of ancestry from distinct populations. An alternative but unlikely explanation of such data is that the parents have ancestries that alternate between the two populations across chromosomes. Thus, considering the genome-wide fraction of each type of unphased local ancestry call in the focal sample (i.e., homozygous for population 0 or 1 or heterozygous for ancestry) allows us to formulate a parsimonious model of the parents’ ancestry fractions. These observations, which may not be fully captured in some other models of parental ancestry, are the key motivation behind PAPI’s binomial model. We note that the approach described below is equivalent to that adopted in another recently developed method to model parental ancestry^28^.

PAPI’s binomial model treats the chromosome transmitted by a given parent *j* ∈ {*A, B*} as a sequence of Bernoulli trials parameterized by the parent’s ancestry proportion—i.e., by the target of inference *p*_*j*_. In fact, a more precise definition of *p*_*j*_ is the fraction of parent *j*’s *transmitted haplotype* contained in tracts of ancestry 0, calculated using Morgan units. This *p*_*j*_ will in general differ from parent *j*’s true ancestry proportion due to recombination and the random assortment of alleles to gametes, and is only equal to the parent’s true ancestry proportion in expectation.

Formally, let ***X*** = ***x***_1_, …, ***x***_*N*_ be the input unphased diploid local ancestry tracts of a focal sample. Each of these tracts is a tuple ***x***_*i*_ = (*l*_*i*_, *a*_*i*_) for *i* ∈ {1, …, *N*} where *l*_*i*_ represents the tract length in Morgans and *a*_*i*_ ∈ {{0, 0}, {0, 1}, {1, 1}} is its unphased local ancestry (Figure 1B). Curly braces in the ancestry component indicate that these values are unordered, since the data are unphased, so the {0, 1} class is consistent with the phased, ordered states (in square brackets) [1, 0] and [0, 1].

The likelihood of the binomial model is:

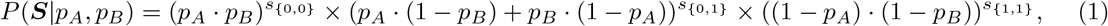

where ***S*** = (*s*_{0,0}_, *s*_{0,1*}*_, *s*_{1,1*}*_) and 1 − *p*_*j*_ is the fraction of parent *j*’s transmitted haplotype contained in tracts of ancestry 1. Let *I* be the indicator function, then *s*_*a*_ =Σ_*i*_ *l*_*i*_ · *I*(*a* = *a*_*i*_) · *τ*, i.e., the total Morgan length of the genome contained in tracts of ancestry type *a*, scaled by a hyperparameter *τ* to account for the effective number of independent markers. This is necessary because the segment lengths are in Morgan units, and the effective number of independent loci in a tract of length 1 Morgan is the number of crossovers that have occurred in that interval since admixture began. In expectation, this number is exactly *t*_*j*_ for the haplotype parent *j* transmitted. Following this reasoning, we set *τ* to a rough population averaged number of generations since admixture. For the African American datasets (both simulated and real) analyzed in this study, we set *τ* = 6, consistent with average estimates of the time since admixture from previous studies in this population^7,29,30^, and with PAPI’s own estimates in the HapMap African ancestry in the Southwest USA (ASW) population of African American samples^31^ (Results).

### Modeling the relationship between admixture time and ancestry switch rates

Crossover placement along a haplotype that was subject to *m* meioses can be modeled as a Poisson process with rate *m* per Morgan. If *m* ≥ 4, this model provides a good fit to intercrossover distances generated under a crossover interference model^26^. However, applying the Poisson model to local ancestry segments is complicated by the presence of within-population *hidden* crossovers. These are historical crossovers that occur at homozygous ancestry positions and, as a result, are not directly observed in local ancestry tracts.

Their presence requires an adjustment to the switch rate parameter—i.e., this rate is not a direct measure of the number of meioses since admixture.

In this subsection we assume that for each parent *j* ∈ {*A, B*}, all the ancestors *t*_*j*_ generations ago are unadmixed and that at least one couple in that generation includes individuals of both ancestries, while the other individuals have the same ancestry (Figure 1A). We refer to this form of pedigree throughout as the *base case*, and it corresponds to a single pulse of admixture, with migrants entering only *t*_*j*_ generations ago. This means that at least one of the ancestors *t*_*j*_ − 1 generations ago is entirely heterozygous for ancestry. We do this to defer consideration of the crossover rate parameter in more complex admixture scenarios—those with multiple pulses of admixture—to later in the text (see “Simulating multiple pulses of admixture on a pedigree”).

The model we adopt approximates pedigree-generated crossovers by Markovian transitions between a pool of haplotypes (Figure 2). The proportion of haplotypes in the pool from population 0 is *p*_*j*_, the parental ancestry proportion. Gravel proposed a population genetic model of this form in the context of pulse admixture events^9^, and noted that the probability of transitioning to a haplotype of a different ancestry is simply the proportion of haplotypes of that ancestry in the population. In particular, the Poisson rates 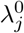 for switching from ancestry 0 to 1, and 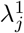 for switching from 1 to 0, are:

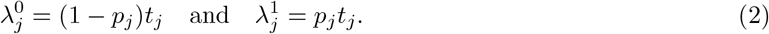

**Figure 2:**
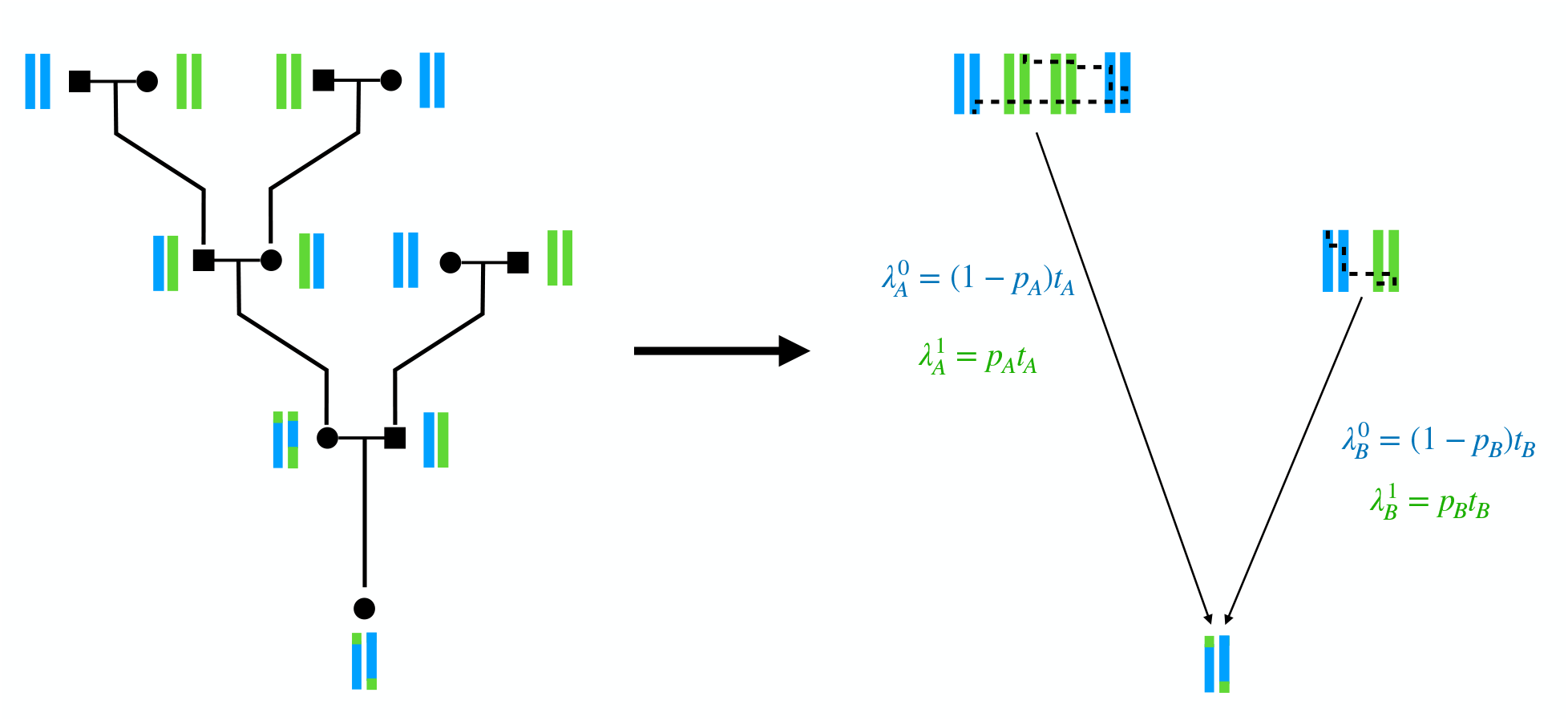
Markovian approximation to an individual’s full pedigree. Example pedigree for an admixed individual (left). Viewing the unadmixed ancestors as a pool of founder haplotypes (right), one for each parent, and further viewing recombinations as a Markovian switching process in this pool motivates the use of a single-pulse migration model to capture the effect of *t*_*j*_ and *p*_*j*_ on the observed between-ancestry Poisson switch rates 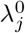 and 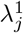.

Thus, to obtain the switch rate out of a given ancestry, the adjustment to the crossover rate *t*_*j*_ is the proportion of haplotypes of the opposite ancestry. Intuitively, one can view these rate parameters 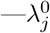 and 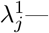 as the expected count of crossovers per Morgan that result in an ancestry switch.

Note the nature of the approximation made here, which is two fold: first, crossovers in the model are no longer constrained to occur within an individual, which ignores long range correlations caused by within-individual meioses^32^. Second, *p*_*j*_ is used to represent the proportion of founder haplotypes of 0 ancestry even though this value—parent *j*’s ancestry proportion—may differ from the fraction of founders that have this ancestry. Despite these caveats, our empirical results suggest that this is an effective approximation (Results).

An important limitation in applying the equations in (2) is that they incorrectly account for the crossovers transmitted by an individual that is entirely heterozygous for ancestry. To illustrate this, consider such a parent *j*, who will have *p*_*j*_ = 0.5. Equation (2) implies that 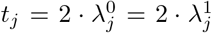, and because every locus in parent *j*’s genome is heterozygous for ancestry, all of *j*’s crossovers will produce ancestry switches. This makes the rate of switches 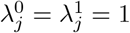 per Morgan, and so, despite including only one meiosis, the equations will suggest that *t*_*j*_ = 2. The issue is that the model implicitly assumes that the ancestry proportions of parent *j*’s two chromosomes are both 0.5 when in fact one is entirely composed of ancestry 0 and the other of ancestry 1. This scenario arises for all ancestors with entirely heterozygous ancestry, and we find that subtracting 1 from the raw estimates is an effective fix no matter how large *t*_*j*_ is^26^ (data not shown). PAPI applies this correction internally by subtracting 1 from the *t*_*j*_ point estimates and from each MCMC sample followed by rounding up negative values to 0.

### A hidden Markov model to jointly infer admixture times and ancestry proportions

PAPI computes the likelihood of the observed unphased local ancestry tracts ***X*** given assignments of the parameters ***θ***, and uses either a Markov chain Monte Carlo (MCMC) or gradient descent method to estimate these parameters (below). The hidden states of the HMM encode phased local ancestry assignments ***z*** = ***z***_1_, ***z***_2_, …, ***z***_*N*_, where ***z***_*i*_ ∈ {[0, 0], [0, 1], [1, 0], [1, 1]} ∀*i*, and the two elements of ***z***_*i*_ correspond to a transmitted chromosome by a given parent; as a matter of convention, we let the first element *z*_*i*,1_ be the chromosome transmitted by parent *A*. Because the phase is unknown, the HMM integrates over all possible phase assignments to compute the likelihood of the unphased data. In order to model the relationship of *p*_*j*_ and *t*_*j*_ to the observed data while accounting for hidden recombinations (above), PAPI internally reprameterizes ***θ*** = (*p*_*A*_, *p*_*B*_, *t*_*A*_, *t*_*B*_) as 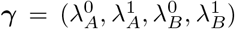 by applying Equation (2) for *j* ∈ {*A, B*}, and the method therefore calculates the likelihood *P* (***X***|***θ***) = *P* (***X***|***γ***). Below, we treat the conditioning on parameters as implicit and omit them from the probabilities.

The HMM models the local ancestry tract lengths as exponentially distributed, as follows from a Poisson crossover model. More specifically, the initial probability is a product of the probabilities of the first observed tracts (for haplotypes *A* and *B*) in ***z***_1_, both of which have length 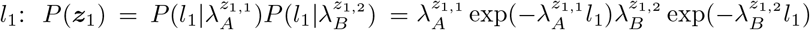. We define the transition probabilities to ensure that tracts that extend beyond the interval they start in have a total probability consistent with their full length. That is, the transition probability *P* (***z***_*i*_|***z***_*i*−1_) depends on whether the ancestry of a haplotype changes between adjacent intervals, and only includes the leading *λ*_*j*_ term in the first interval where haplotype *j* has switched ancestries.

We express these as:

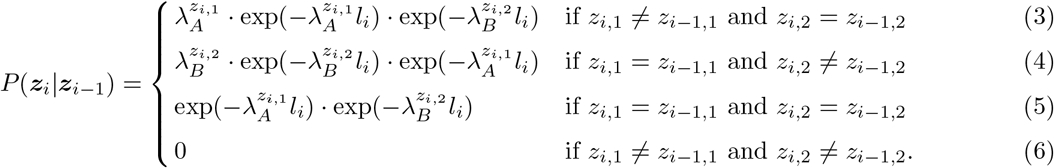

For example, in Equation (3), parent *B*’s transmitted chromosome does not change ancestry (*z*_*i*,2_ = *z*_*i*−1,2_), and the transition probability incorporates the 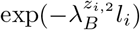 term to reflect this. The total probability of this tract from its origin in a given interval *k* to its termination at the end of some interval *l* then is 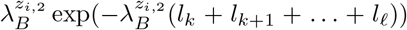—which is identical to the probability of observing a tract of length (*l*_*k*_ + *l*_*k*+1_ + …+ *l*_**l**_). By contrast, for Equation 3 to apply, haplotype *A* would have changed ancestries, and the formula includes a leading 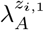 term here where the tract begins. Other cases follow this same pattern, but the final case (in Equation 6) requires two crossovers to occur at the same position (one on each haplotype). This is prohibited by the model, and the corresponding transition probability is set to 0. If the observed data implies such a transition and PAPI is run without an error model (below), it will throw an error. Although in our analyses such transitions never occurred in the HAPMIX local ancestry tracts, a user can address any such cases by pre-processing them to introduce a small tract that switches the ancestry of only one haplotype between the two disallowed tracts.

The above formulation is somewhat non-standard in that the initial and transition probabilities depend on the tract lengths, which are part of the observed data. An alternative might put the exponential terms— which are calculated from the tract lengths—into the emission probabilities, leaving only the leading *λ* terms in the transition probabilities. However, doing this yields transition probabilities that are typically greater than one and obscures the conceptual underpinnings of the model—that of a Poisson processes. The current formulation can be thought of as implicitly combining more standard transition and emission terms in the transition probabilities. We therefore set the emission probabilities in this formulation to 1 if the hidden state is a phased version of the observed local ancestry, and 0 if not. We describe an optional error model that allows for mismatched hidden and observed ancestry states in the next subsection.

We treat the final tract of a chromosome as partially observed, as the position of the next crossover is unknown and would land somewhere beyond the chromosome endpoint. Such tracts should contribute only an exponential term to the likelihood, with the leading *λ*_*j*_ term omitted—corresponding to the probability of a tract of length greater than that observed. PAPI accomplishes this by setting the probability of transitioning from the final tract to a special end state Ω as: 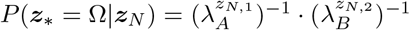. These inverse terms act to cancel out the 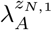 and 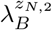 terms that were introduced when the tracts with ancestry *z*_*N*,1_ and *z*_*N*,2_ were first transitioned to, leaving only exponential terms, as desired.

With the HMM’s initial, transition, and emission probabilities specified, PAPI computes the likelihood of the observed data ***X*** given ***θ*** by marginalizing over all possible hidden state paths *P* (**x**|***θ***) =Σ_**z**_ *P* (**x**|***θ*, z**) using the forward algorithm. The model treats each chromosome as independent given ***θ***, so the total likelihood is the product of each chromosome’s likelihood. Note that the fact that parent labels are arbitrary and interchangeable introduces non-identifiability into the model. In particular, ***θ*** = (*p*_1_, *p*_2_, *t*_1_, *t*_2_) and ***θ***^*^ = (*p*_2_, *p*_1_, *t*_2_, *t*_1_) have the same likelihood and only produce swapped parent labels. When run in the MCMC inference mode (see below), PAPI handles this non-identifiability by restricting the parameter space to points such that *p*_*A*_ *> p*_*B*_. When run in the gradient descent inference mode (below), there is no need to restrict the space, as PAPI will find a mode of the posterior distribution, and the presence of more than one equivalent mode does not hamper the inference.

The above HMM is similar to that of ANCESTOR^14^, another approach for inferring parent ancestry proportions. Both methods apply to unphased local ancestry calls and model the phase of those calls in the hidden states. A point of departure is that PAPI parameterizes the model explicitly in terms of ***θ*** (although it still performs the underlying calculations in terms of ***γ***), which affords it two advantages. First, by modeling hidden crossovers (above), PAPI can produce direct estimates of *t*_*A*_ and *t*_*B*_ rather than only *λ*_*A*_ and *λ*_*B*_. Second, PAPI’s HMM uses more of the available information for inferring (*p*_*A*_, *p*_*B*_) by incorporating these parameters into the transition probabilities (via *λ*_*A*_ and *λ*_*B*_), thus producing more accurate estimates of (*p*_*A*_, *p*_*B*_) across a range of scenarios (Results). While the (*p*_*A*_, *p*_*B*_) estimates from PAPI’s HMM are in general more accurate than those of ANCESTOR, for the most accurate *p*_*j*_ estimates, PAPI’s binomial component is important (especially when |*p*_*A*_ − *p*_*B*_| is large), and PAPI performs best when run with the HMM and binomial component together using the full model (Results).

### Handling erroneous tracts

Despite being immune to switch errors, unphased local ancestry segments are still subject to erroneous calls that have the potential to bias the inference of *t*_*j*_. We address this by providing an optional error model that has a non-zero probability of emitting unphased local ancestry tracts with a different ancestry than that of the hidden state. Using the congruence operator, we say ***z*** ≅ *a* if the ordered hidden state *z* has the same ancestry values as those in the unordered observed local ancestry *a*; e.g., [1, 0] ≅ {0, 1} and [0, 0] ≇ {0, 1}. Using this notation, the emission probabilities of the error model are:

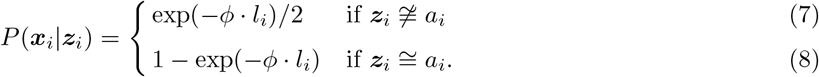

Here *ϕ* is a hyperparameter that we set heuristically so that when ***z***_*i*_ ≇ *a*_*i*_, *P* (***x***_*i*_|***z***_*i*_) decreases by a factor of 1*/*2 for every centiMorgan unit increase in *l*_*i*_, so *ϕ* = 100 · ln 2. We divide the exponential term in (7) by 1/2 because there are three unordered local ancestry calls and therefore two ways to select *a*_*i*_ such that ***z***_*i*_ ≇ *a*_*i*_. The exponential form of these equations leads to higher emission probabilities for incongruous tracts that are small. Another way to think about this is that the model applies a penalty for treating larger tracts as erroneous, and that the penalty decreases exponentially with decreasing tract size. So if the HMM encounters a sufficiently small tract where *a*_*i*_ differs from *a*_*i*−1_ and *a*_*i*+1_, it may favor the incongruous hidden state (i.e, where ***z***_*i*_ ≇ *a*_*i*_ but ***z***_*i*_ = ***z***_*i*−1_ = ***z***_*i*+1_) in order to avoid ancestry state switches over a small distance (see Equations (3) and (4)) while incurring the incongruous tract penalty from Equation (7).

### Combining component models and performing inference

As noted earlier, PAPI can combine the output signals from its two component models—the binomial model and the HMM—which improves its inference of ***θ*** compared to either model alone (Results). Notably, both models operate on the same data ***X***, which renders their likelihoods non-independent, but the HMM analyzes the unphased local ancestry calls ***X*** directly, while the binomial model considers the summary statistics in ***S***. Despite this non-independence, PAPI’s default setting combines them to form a composite likelihood, treating *P* (***X, S***|***θ***) ≈ *P* (***X***|***θ***)*P* (***S***|***θ***). Furthermore, PAPI uses a Bayesian model, computing the following posterior based on the composite likelihood (which we refer to as the *full* model):

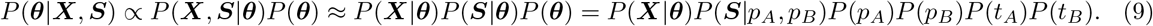

Here we make explicit the assumption that all the model parameters (in ***θ***) are independent, and note that the binomial likelihood *P* (***S***|***θ***) = *P* (***S***|*p*_*A*_, *p*_*B*_) since that model only considers (*p*_*A*_, *p*_*B*_). We use uniform priors on the parameters, with *p*_*A*_, *p*_*B*_ ∼ *U* (0, 1) and *t*_*A*_, *t*_*B*_ ∼ *U* (0, 25).

PAPI uses one of two methods to perform inference, as specified by the user. By default, it performs gradient descent using the constrained optimization algorithm L-BFGS-B (implemented in the scipy-optimize Python package). This yields a point estimate corresponding to a local maximum of the likelihood function, and we refer to this as a GD estimate. The empirical results suggest that these estimates, despite being local maxima, are reliable in practice. The second inference method uses MCMC in order to draw from the posterior distribution—specifically performing slice sampling (via the PyMC3 package). In this setting, PAPI performs 500 burn-in iterations followed by 1,000 MCMC draws across 4 independent chains per individual (6,000 samples total) and outputs a 90% credible interval for each of the parameters.

The GD inference method is quite fast, whereas the MCMC approach requires larger compute times. In light of timing and performance (see Results), we recommend running PAPI using the full model (Equation 9) in the GD inference mode. It is also possible to run the HMM or binomial model alone, i.e., to analyze *P* (***θ***|***X***) or *P* (***θ***|***S***), respectively. (In these settings, PAPI does not leverage a composite likelihood.) Note that all three models supply *p*_*j*_ estimates, and a user can run the binomial model in MCMC mode if posterior distributions of *p*_*j*_ alone are required (which we recommend, as the MCMC runs are much faster with the binomial model alone—Results).

### Simulating realistic admixed genotypes

Perhaps the most realistic way to simulate genetic data for an admixed individual would be to explicitly model their pedigree, with its unadmixed founder ancestors, and simulate a meiosis for each of the ancestors. This would appropriately constrain crossovers to occur only between each ancestor’s two haplotypes, and can incorporate crossover interference modeling^26^. However, doing this requires the use of data from 2^*T*^ founders, where *T* is the number of ancestral generations in the pedigree, thus consuming large data resources from unadmixed individuals to simulate only a single admixed individual. Furthermore, some unadmixed samples must be held out of the simulation for use in the panels required by the local ancestry inference software.

We adopted a three-step hybrid simulation scheme that explicitly accounts for pedigree-based transmissions while using far fewer unadmixed samples (Supplementary Figure 1). First, we ran Ped-sim^26^ to simulate individuals from a specified pedigree structure, similar to the above approach, but without providing input genetic data, and thus obtained no output genotypes at this stage. We used a sex-specific genetic map^33^, a Poisson crossover model, and ran Ped-sim in a mode (using the `--bp` option) that outputs the founder haplotype id that the focal admixed individual carries at each position with crossover break points indicated. Rather than using the resulting break point file as is, with its 2^*T*^ unique haplotypes, in step two we modified it by mapping each founder haplotype to one of the two ancestral population labels according to the desired value of *E*[(*p*_*A*_, *p*_*B*_)] as well as (*t*_*A*_, *t*_*B*_) (see below). Finally, we used this population-only break point file as input to admix-simu^34^, which also takes in genetic data corresponding to unadmixed founder haplotypes from each population. Admix-simu treats the founder haplotypes as a pool, and samples haplotype segments based on the population labels and break points contained in the break point file. More specifically, at each break point, the method randomly samples a new haplotype (of the appropriate population) from the pool, creating non-overlapping sampling paths (Supplementary Figure 1). This interrupts the linkage disequilibritum (LD) on either side of a break point as in a real crossover and approximates the full pedigree model. After sampling all segments of the target admixed individuals’ haplotypes, admix-simu outputs the final haplotypes, which we then grouped in pairs to form individual diploid genotypes.

We simulated pedigrees of the base case form introduced earlier (see “Modeling the relationship between admixture time and ancestry switch rates”), assigning the founders in generations (*t*_*A*_, *t*_*B*_) to different populations according to the proportions given by the expectation *E*[(*p*_*A*_, *p*_*B*_)]. This expectation will in general differ from the realized values (*p*_*A*_, *p*_*B*_) (in the transmitted parental haplotypes of the focal individual) that we used to calculate deviation statistics (see “Computing measures of accuracy and bias”). Further, the range of *E*[(*p*_*A*_, *p*_*B*_)] values is constrained by the number of founder individuals in the pedigree, which depends on (*t*_*A*_, *t*_*B*_) (e.g., it is impossible to simulate *E*[(*p*_*A*_, *p*_*B*_)] = (0.1, 0.1) when (*t*_*A*_, *t*_*B*_) = (2, 2) as the number of founder haplotypes is too small). Therefore, we assigned the founder ancestries to approximate the expected proportions as closely as possible.

We used data from the HapMap CEU and YRI populations as the input unadmixed founder haplotypes for simulating. In order to provide large unadmixed panels to the local ancestry inference software and ensure these are unrelated to the target simulated individuals, we first excluded children from the HapMap CEU and YRI populations and split the remaining individuals (trio parents, duo parents and unrelated samples) into four sets of 22 individuals for the two populations. We then used the pipeline described above to simulate admixed individuals in four separate runs, each using one of the four sets of simulation founders as input to admix-simu, and with each assigned a different *E*[(*p*_*A*_, *p*_*B*_)] value: (0.25, 0.75), (0.4, 0.6), (0.25, 0.5) or (0.5, 0.5). We repeated this procedure (using the same four founder datasets) for each of the desired settings of (*t*_*A*_, *t*_*B*_) (see Results for these values). This produces four sets of simulated samples for every (*t*_*A*_, *t*_*B*_) that are unrelated to the other three sets of unadmixed samples. Doing this allows us to utilize the three sets of unadmixed samples (66 individuals in total) as panels in HAPMIX when analyzing the corresponding simulated individuals, thus leveraging all the available data for simulating while still maintaining sizeable panels for local ancestry inference.

### Simulating multiple pulses of admixture on a pedigree

The base case pedigree only includes a subset of the admixture histories that are possible, as unadmixed ancestors from distinct populations will in general occur at more than one generation in an admixed individual’s pedigree. We sought to ask what value of *t*_*j*_ our pulse-migration model (Equation (2)) will estimate in more general cases.

To characterize this, we simulated admixed haplotypes that have two admixture time parameters, 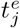 and 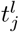 which are, respectively, the earlier and later generations 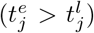 in which couples containing unadmixed individuals of different ancestries occur (all other individuals are unadmixed of the majority ancestry). To perform these simulations, we used a base case pedigree for 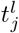 with a desired setting of *E*[*p*_*j*_] and generated new pedigrees by replacing lineages—starting from the left-most lineage and working to the right—by lineages with a time since admixture of 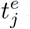 and with the same *E*[*p*_*j*_]. We use the parameter *q* to define the proportion of all the couples in the 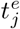 generation composed of individuals of different ancestries. By this definition, the range of *q* depends on *E*[*p*_*j*_], as the maximum fraction of couples containing unadmixed individuals of different ancestries is 2*E*[*p*_*j*_], so when *E*[*p*_*j*_] = 0.5, *q* ranges between 0 and 1 and when *E*[*p*_*j*_] = 0.25, *q*’s maximum value is 0.5.

Note that for this analysis we used only the exact local ancestry tracts derived from the Ped-sim break point positions. We represent the tracts lengths from population 0 as 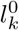 for *k* ∈ {1, …, *N*^0^} and from population 1 as 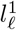 for *l* ∈ {1, …, *N*^1^}. From these we estimated the Poisson rates using the maximum likelihood estimator for the 22 finite autosomes^26^ as 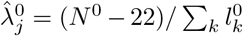 and 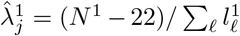. Finally, from these same data we also estimated 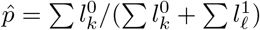.

In order to calculate the value of *t*_*j*_ the model in Equation (2) infers, we reasoned that 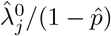 and 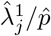 each give a measure of *t*_*j*_, and we compute a weighted average of these two estimators. Specifically, since the total Morgan fraction of the genome that fall in tracts of ancestry 0 and 1 are 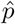 and 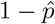 respectively, we calculated:

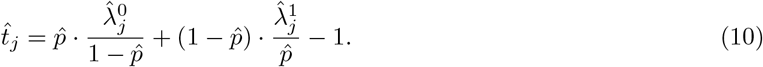

Here, as in PAPI, we subtract 1 for reasons described earlier (see “Modeling the relationship between admixture time and ancestry switch rates”).

### Handling local ancestry calls as input to PAPI

PAPI reads local ancestry calls produced by both HAPMIX and LAMP-LD. HAPMIX outputs posterior probabilities of the three possible unphased diploid local ancestry states at every marker, which PAPI converts into an internal representation of fixed (i.e., non-probabilistic) local ancestry calls. This conversion works by taking the maximally probable state at each marker. In contrast, LAMP-LD outputs fixed local ancestry calls and PAPI converts them directly into an internal representation. In simulated data, we found that LAMP-LD is prone to producing erroneous short segments, leading to an upward bias in admixture time inference. To address this, we filtered out LAMP-LD tracts shorter than 1 cM, which is effective at correcting this bias (results not shown). Note that filtering out tracts can introduce state transitions prohibited by the HMM (see “A hidden Markov model to jointly infer admixture times and ancestry proportions”). In order to avoid this, we filtered out only those tracts that are flanked by tracts of identical ancestry state. The filtering process consists of deleting the target tract and merging the flanking tracts to fill in the missing space. We applied this filter to all the LAMP-LD-based analyses.

### Computing measures of accuracy and bias

The *A* and *B* parent labels in PAPI’s 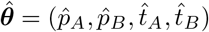 estimates separate the inferred time and ancestry proportions of one parent from the other. However, PAPI does not infer the sexes of these parents, and in order to compute measures of accuracy and bias in our simulations, it is necessary to map these labels to specific parents. To do this, we choose the assignment that minimizes the sum of squared errors. Let *p*_*max*_ and *p*_*min*_ be the ancestry proportions of the parent with the larger and smaller estimates, respectively. Then, representing the estimated labels by an ordered vector ***v*** ∈ {(*A, B*), (*B, A*)}, we minimize 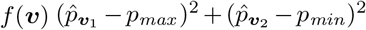 and use ***v*** ^**\***^= arg min_***v***_ *f* (***v***) as the mapping between estimated and true parent labels.

In Results, we report the mean absolute deviation of the estimated parameters from the truth, which serves as a measure of accuracy, computing this as

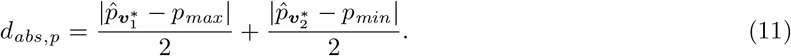

We also report a measure that uses exact rather than absolute differences, which serves as a measure of bias. This statistic is a vector of two numbers corresponding to the bias of each parent’s ancestry proportion individually:

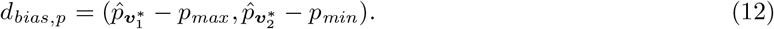

We also compute analogous metrics for the 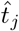 estimates, denoted *d*_*abs,t*_ and *d*_*bias,t*_.

### Filtering and processing the PAGE data

We used PAPI to analyze African American individuals from the BioMe Biobank subset of the Population Architecture using Genomics and Epidemiology (PAGE II; dbGaP:phs000925.v1.p1) study^27^. In order to identify African American samples, we merged all the individuals (*N* =12,754) with the HapMap CEU and YRI trio parents—the latter providing known population labels—and ran ADMIXTURE v1.3^35^ with *K* = 5. We analyzed the estimated ancestry proportions for the HapMap individuals to determine which of the *K* populations corresponded to African and European ancestry. We then selected PAGE individuals that met each of the following three criteria as subjects for our study: (1) has ≥5% African ancestry; (2) has ≥5% European ancestry; and (3) the sum of these two ancestries is ≥99.5%. This identified a total of 5,786 subjects for our analysis.

Because the number of SNPs that intersect between the PAGE and HapMap III datasets is small, we used the YRI and CEU trio parents from the 1,000 Genomes Project^36^ (1000G) data as panels for performing local ancestry inference in the PAGE subjects. To get a common set of SNPs between 1000G and PAGE, we first filtered out multiallelic and indel variants from both datasets. Next, in order to prevent any strand inconsistencies in the allele encodings of the two datasets, we ran PLINK v2.00a2.3LM with the `--ref-from-fa` option to recode the PAGE alleles to the forward strand of the GRCh37 reference genome (the 1000G data is already encoded in this manner). This may not correctly assign alleles at A/T and C/G SNPs, and we filtered these SNPs later (below). Next we applied the “composite filter” provided in the quality control report distributed with the PAGE data, which includes filters for Hardy-Weinberg equilibrium, discordant calls in duplicated samples, and Mendelian errors, among others. In order to identify any remaining incorrectly coded alleles, we applied a chi-squared test for allele frequency differences between two groups of subjects—the 5,786 PAGE individuals and a custom subset of 1000G composed of 80 YRI and 20 CEU individuals. Markers in the PAGE data and the custom dataset are expected to be have similar allele frequencies, and those with statistically significant differences indicate a potential mismatch in the strand encoding. We filtered all markers with |*z*| *>* 4 from both datasets. Finally, we removed all A/T and C/G SNPs, as allele encoding mismatches in these markers (e.g., at high minor allele frequency variants) may not be detected by any of the analyses above. All of these steps left 409,095 markers that overlapped between PAGE and the 1000G data, and we used these to perform local ancestry inference via HAPMIX on each PAGE sample with the 1000G data as panels.

## Results

We validated PAPI’s approach to deconvolving parental transmissions within unphased diploid local ancestry tracts of a given individual using simulated and real data. Our first analysis contrasts the accuracy of PAPI’s estimates using each of its models and inference modes. Following this, we compare PAPI’s parameter estimates of (*p*_*A*_, *p*_*B*_) with those of ANCESTOR (which does not estimate (*t*_*A*_, *t*_*B*_)) under a number of simulated scenarios, finding that PAPI reliably infers these parent ancestry fractions. Next, we turn to more deeply examining the quality of PAPI’s (*t*_*A*_, *t*_*B*_) estimates using both inferred local ancestry tracts and exact tracts from our simulation pipeline. For these analyses, we considered a range of admixture histories, including the base case and more complex pedigrees (Methods).

Following these efforts, we applied PAPI to a large dataset of African Americans from the PAGE study. By leveraging PAPI’s power to accurately infer parental ancestry proportions, we examined the relationship of couples’ ancestries to one another and find strong evidence of non-random mating, as described below.

### Evaluating the performance of PAPI’s component models

We simulated the genotypes of admixed individuals under three scenarios: (1) *t*_*A*_ *> t*_*B*_, with *t*_*A*_ = 9 and *t*_*B*_ ∈ [2, 8]; (2) *t*_*A*_ = *t*_*B*_ ∈ [2, 9]; and (3) (*t*_*A*_, *t*_*B*_) ∈ {(0, 0), (1, 1)} (see Methods for simulation details). For every (*t*_*A*_, *t*_*B*_) in scenarios (1) and (2), we simulated 22 admixed individuals for each of *E*[(*p*_*A*_, *p*_*B*_)] ∈ {(0.5, 0.5), (0.25, 0.5), (0.4, 0.6), (0.25, 0.75)} for a total of 88 individuals per scenario. In scenario (3), the small numbers of generations correspond to limited ranges of parent ancestry proportions; as such, when (*t*_*A*_, *t*_*B*_) = (0, 0), we simulated 22 admixed individuals for each of *E*[(*p*_*A*_, *p*_*B*_)] ∈ {(1, 0), (0, 0), (0, 1), (1, 1)}, and when (*t*_*A*_, *t*_*B*_) = (1, 1), we simulated 22 admixed individuals for each of *E*[(*p*_*A*_, *p*_*B*_)] ∈ {(0.5, 0.5), (0, 0.5)} for a total of 132 individuals. Note that here and elsewhere in Results, *p*_*j*_ corresponds to African ancestry proportions. As described in Methods, our simulation pipeline assigns unadmixed founders in a pedigree to population labels to match as closely as possible the desired parent ancestry proportions. Because of random assortment and recombination, the realized (*p*_*A*_, *p*_*B*_) vary, and in general these founder ancestry assignments only give the expectation *E*[(*p*_*A*_, *p*_*B*_)] (or more precisely, the closest possible values to this expectation). However we used the realized values of (*p*_*A*_, *p*_*B*_) for computing deviation metrics *d*_*abs,p*_ and *d*_*bias,p*_.

PAPI can estimate (*p*_*A*_, *p*_*B*_) using any of its three models: the binomial, the HMM, or the full model (a composite of the binomial and the HMM; Methods). We ran all three models on the scenario (1) simulated individuals (Supplementary Figure 2), and, averaged across all data points taken together, the GD estimates of (*p*_*A*_, *p*_*B*_) under the full model are quite reliable with *d*_*abs,p*_ = 0.043. This is the best performing GD-based model overall with an absolute deviance 6.52% and 14.0% smaller than those of the HMM (*d*_*abs,p*_ = 0.046) and binomial model (*d*_*abs,p*_ = 0.050), respectively. The same trend holds for the MCMC estimates, albeit with lower relative differences among the models: averaged across all scenario (1) data points, the binomial, HMM, and full models achieve *d*_*abs,p*_ values of 0.042, 0.041, and 0.040, respectively. The differences between the models are most pronounced when *E*[(*p*_*A*_, *p*_*B*_)] = (0.5, 0.5); in particular, using the GD inference mode for this setting, the *d*_*abs,p*_ statistics for the binomial, HMM, and full models are, respectively, 0.061, 0.054, and 0.053; in turn, using MCMC inference, the corresponding *d*_*abs,p*_ measures are 0.061, 0.054, and 0.049, respectively. Note that while the MCMC (*p*_*A*_, *p*_*B*_) estimates are more accurate than the corresponding GD estimates overall, this method’s increased runtime and loss of accuracy in (*t*_*A*_, *t*_*B*_) estimation (below) make less useful when a user also seeks these time estimates.

Considering PAPI’s (*t*_*A*_, *t*_*B*_) estimates, we measured the Pearson correlation *R* of the inferred *t*_*B*_ estimates with the true simulated *t*_*B*_ values in scenario (1) data (*t*_*A*_ = 9 is held fixed in scenario (1), and we ignore it in the correlation analysis). Here, PAPI’s GD estimates under the full model yield *R* = 0.76, representing a modest improvement compared to the HMM (*R* = 0.71). By contrast, the MCMC results in this case show weaker performance, with the full model yielding *R* = 0.678 and the HMM model alone obtaining *R* = 0.681.

We also benchmarked PAPI’s runtime for every combination of options (inference mode and model) on compute nodes with Intel Xeon E5 4620 processors, utilizing 4 GB of RAM (Supplementary Table 1). We analyzed the 88 individuals simulated under scenario (2) with *t*_*A*_ = *t*_*B*_ = 5, one individual per PAPI run. The average wall clock time for the full model using the MCMC mode is 8.97 hours per individual compared to 23.9 seconds using the GD mode, making the GD mode 1,346× faster than MCMC. The MCMC and GD runtimes for the full model are, somewhat counter-intuitively, slightly shorter than those for the HMM model (6.00% and 9.20% shorter, respectively), while runtimes with the error model option turned on are slightly longer (for the HMM model, 3.29% and 8.75% longer for the MCMC and GD options, respectively). Finally, the binomial model runtimes are quite fast, with the MCMC mode completing in 5.38 minutes and GD mode requiring only 7.3 seconds.

In light of these results, we recommend running PAPI with the full model and using the GD inference option in most cases. These are PAPI’s default settings, and unless otherwise stated, we used these options to produce the rest of the results. A notable exception to this recommendation is when a user seeks only (*p*_*A*_, *p*_*B*_) estimates; in this case the MCMC runtimes for the binomial model are relatively fast, and while this mode provides the least accurate of all MCMC estimates, they are more accurate than the best GD estimates, making the speed and accuracy trade-off worthwhile.

### Accuracy of parent ancestry estimates across admixture scenarios

To evaluate PAPI’s performance in the context of other methods, we compared its inferred parental ancestry proportions with those of ANCESTOR across all three simulation scenarios described above. Figure 3 shows *d*_*abs,p*_ averaged over all scenario (1) and (2) cases (Figure 3A), as well as representative cases from each scenario (Figure 3B and 3C). Overall, PAPI achieves *d*_*abs,p*_ = 0.047 on average for all scenario (1) and (2) cases compared to *d*_*abs,p*_ = 0.087 from ANCESTOR (representing a 46% improvement). PAPI’s performance increases as the difference |*p*_*A*_ − *p*_*B*_| grows; for example, when *E*[(*p*_*A*_, *p*_*B*_)] = (0.25, 0.75), PAPI’s absolute deviance (averaged over admixture times) is 0.032, which is 75.6% lower than ANCESTOR’s corresponding value of *d*_*abs,p*_ = 0.131. We examined the relative performance of PAPI and ANCESTOR in every setting of (*t*_*A*_, *t*_*B*_) from scenarios (1) and (2) and find that PAPI outperforms ANCESTOR in the majority of cases, with a handful of exceptions when *E*[(*p*_*A*_, *p*_*B*_)] = (0.5, 0.5) (see Supplementary Figures 3 and 4).

**Figure 3:**
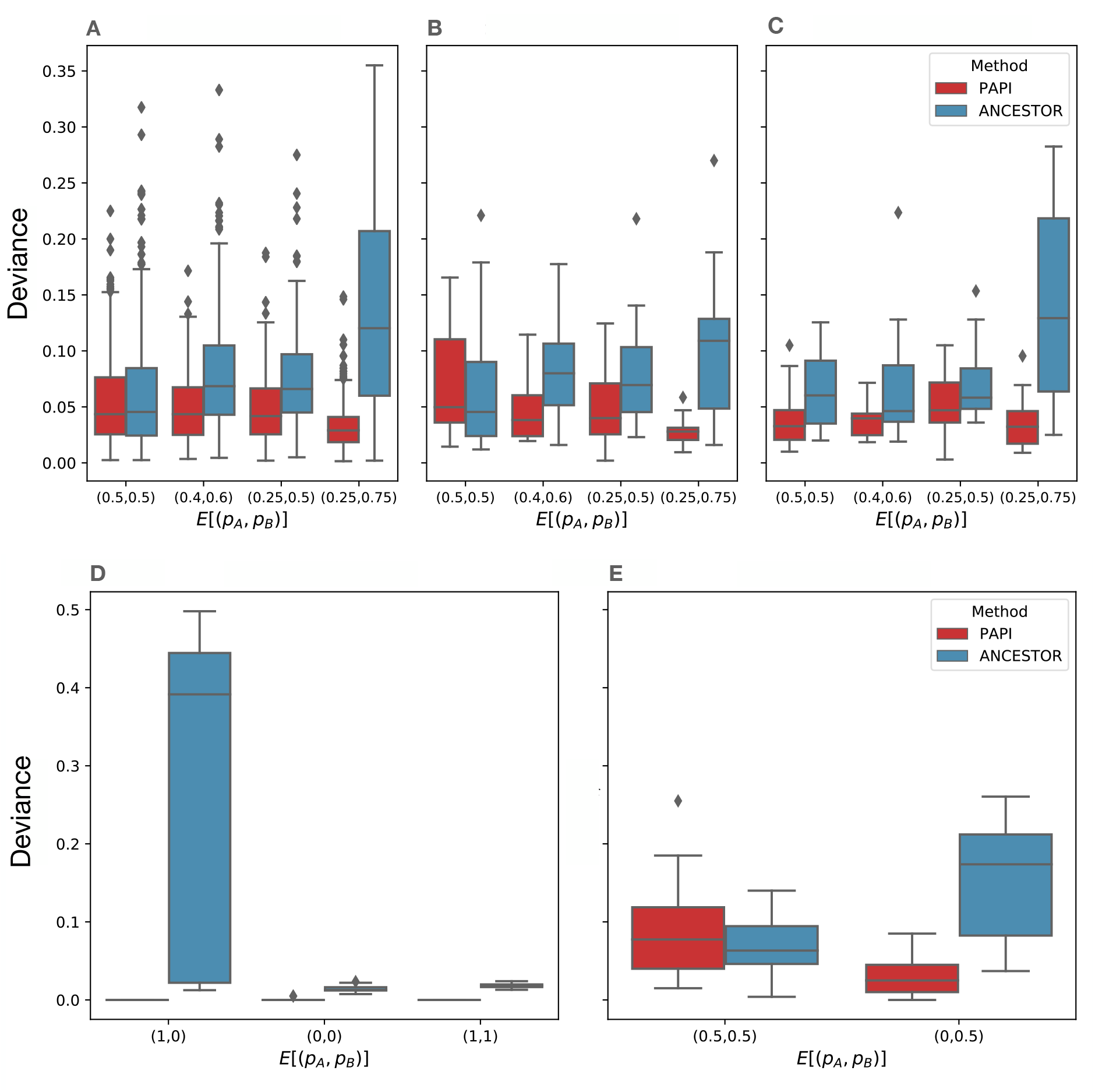
Deviance statistics for inferred parent ancestry proportions (*p*_*A*_, *p*_*B*_) from ANCESTOR and PAPI. (A) Average absolute deviances (*d*_*abs,p*_) on all scenario (1) and (2) data. (B–C) Absolute deviances for data points for (B) *t*_*A*_ = *t*_*B*_ = 5 and (C) *t*_*A*_ = 9 and *t*_*B*_ = 5. (D–E) Average absolute deviance on all scenario (3) data. (D) PAPI estimates the ground truth near-perfectly when *E*[(*p*_*A*_, *p*_*B*_)] ∈ {(1, 0), (0, 0), (1, 1)} and (E) ANCESTOR outperforms PAPI when *E*[(*p*_*A*_, *p*_*B*_)] = (0.5, 0.5) but PAPI’s deviance is low when *E*[(*p*_*A*_, *p*_*B*_)] = (0, 0.5)

PAPI’s parent ancestry estimates have little bias, with average *d*_*bias,p*_ = (0.0085, −0.0078) in scenario (1) samples and *d*_*bias,p*_ = (0.0096, −0.010) in scenario (2) samples. The corresponding values for ANCESTOR are higher at *d*_*bias,p*_ = (−0.056, 0.022) and *d*_*bias,p*_ = (−0.047, 0.035) in scenarios (1) and (2), respectively. Furthermore, the standard deviations of PAPI’s *d*_*abs,p*_ estimates are low at 0.035 in scenario (1) and 0.029 in scenario (2) samples, with corresponding values for ANCESTOR being 0.064 and 0.065, respectively.

Turing to the scenario (3) simulated data, in cases where both parents of the focal sample are unadmixed and from different populations (*E*[(*p*_*A*_, *p*_*B*_)] = (1, 0) or *E*[(*p*_*A*_, *p*_*B*_)] = (0, 1), so the focal sample is heterozygous for ancestry genome-wide), PAPI infers the ground truth with near-perfect accuracy (*d*_*abs,p*_ = 0.0018). In turn, ANCESTOR has fairly high deviance in these cases, averaging *d*_*abs,p*_ = 0.277, with a standard deviation of 0.195 (Figure 3D). When all four grandparents are unadmixed with each couple containing individuals of each ancestry (*t*_*A*_ = *t*_*B*_ = 1 and *E*[(*p*_*A*_, *p*_*B*_)] = (0.5, 0.5)), PAPI is less accurate than ANCESTOR, with the two methods having *d*_*abs,p*_ values of 0.089 and 0.067, respectively (Figure 3E). Even so, when exactly one parent is unadmixed (*t*_*A*_ = *t*_*B*_ = 1 and *E*[(*p*_*A*_, *p*_*B*_)] = (0, 0.5)), PAPI achieves *d*_*abs,p*_ = 0.030, which is considerably better than ANCESTOR’s *d*_*abs,p*_ = 0.150 (Figure 3E).

Finally, we validated PAPI’s parent ancestry estimates in real data by running it on the offspring of all 10 African American trios in the HapMap ASW population. The availability of parental data provides a direct measure of the ground truth via those parents’ local ancestries (which we inferred using HAPMIX). As in the simulated data, we assigned the estimated parent labels to the real parents by minimizing the sum of squared deviances (Methods). Figure 4A shows the estimated and true parent ancestry proportions of the trio children; overall, the inferred values are quite accurate and have fairly low bias, with average *d*_*abs,p*_ = 0.057 and *d*_*bias,p*_ = (0.0272, −0.0263).

**Figure 4:**
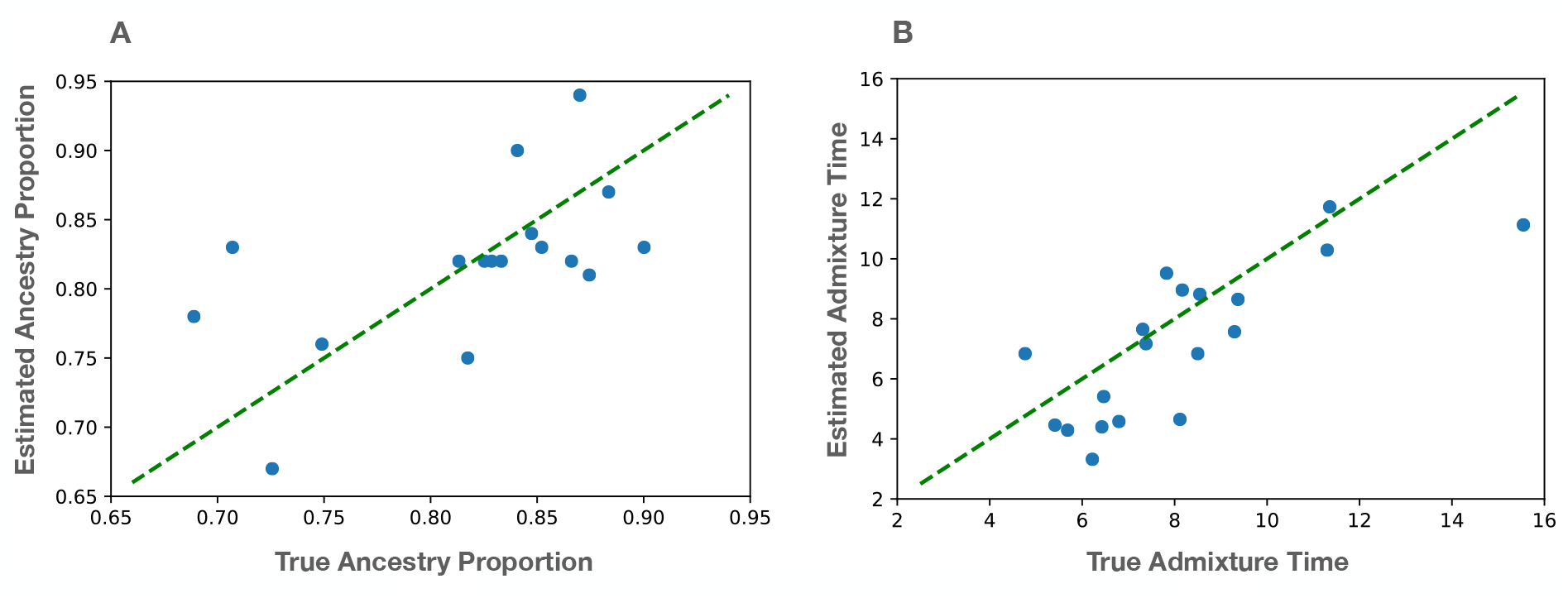
PAPI estimates for the HapMap ASW trio children. Scatter plots of (A) the estimated versus true parent ancestry proportions *p*_*j*_, and (B) the estimated versus true admixture times *t*_*j*_.

### Accuracy of time since admixture estimates

We used PAPI to estimate (*t*_*A*_, *t*_*B*_) in data simulated under scenarios (1) and (2) (above) and found that it reliably recovers the truth under both scenarios (Figure 5A,B). In scenario (1) data, the PAPI estimates for the two parents are well separated, with the difference between the two estimates proportional to the true difference. As noted earlier, there is a high correlation between *t*_*B*_ and the corresponding PAPI estimate (*R* = 0.76), and a more moderate one between the inferred branch differences 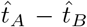 and the truth (*R* = 0.53). PAPI achieves similar quantitative results in the scenario (2) data, where the correlation between its estimates and the truth is *R* = 0.79 (calculated using both the *t*_*A*_ and *t*_*B*_ data points).

**Figure 5:**
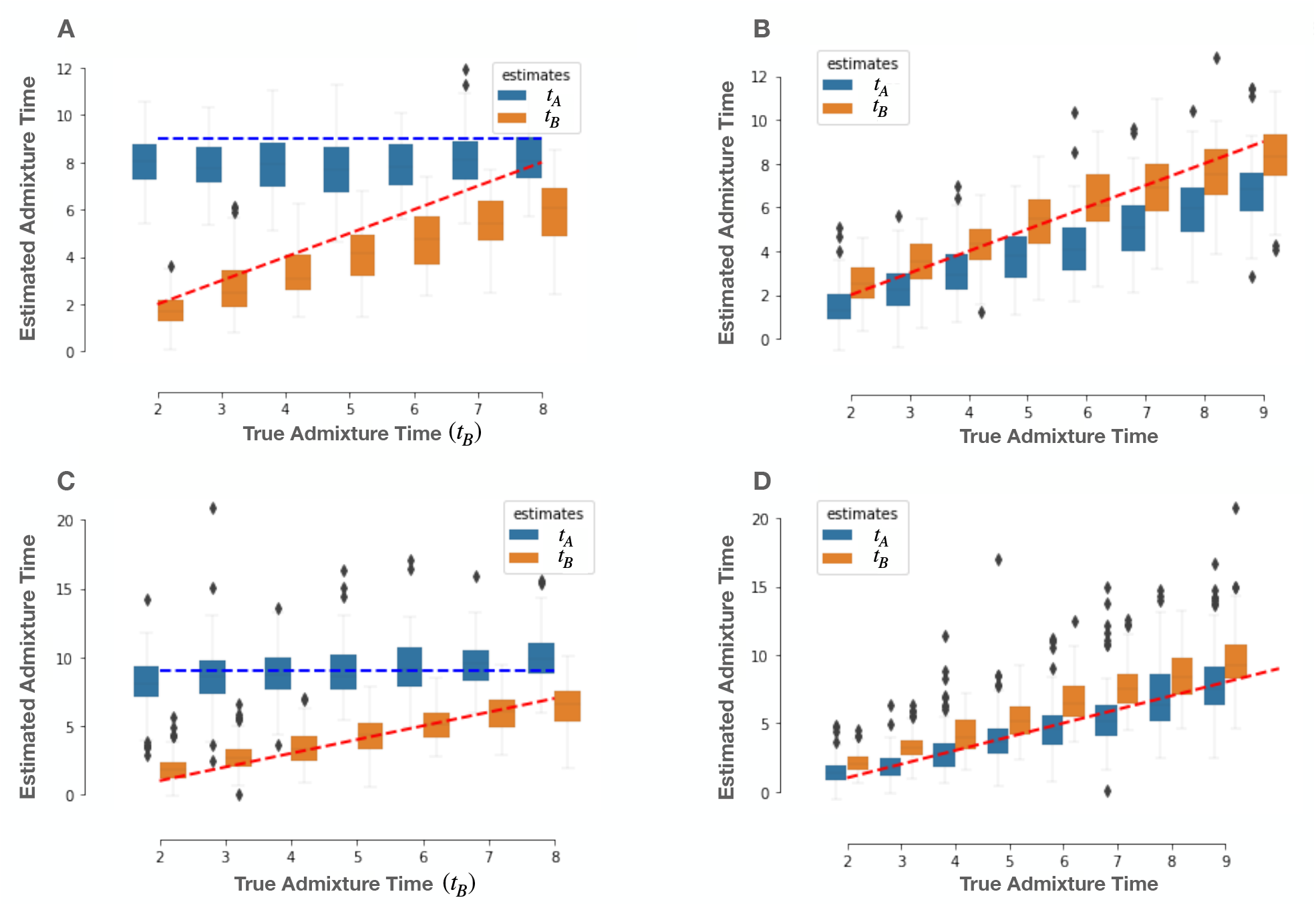
PAPI’s estimated time since admixture in simulated individuals. (A–B) Box plots of estimated times based on HAPMIX local ancestry tracts for (A) scenario (1) samples (where *t*_*A*_ *> t*_*B*_) and (B) scenario (2) data (where *t*_*A*_ = *t*_*B*_). (C–D) Estimated times based on exact local ancestry tracts for (C) scenario (1) and (D) scenario (2) individuals. In panels (A) and (C), the x-axis gives the true *t*_*B*_ since *t*_*A*_ is held fixed in these cases.

PAPI’s estimates in these simulated data have an overall downward bias, with mean *d*_*bias,t*_ = (−1.03, −0.986) for scenario (1) and *d*_*bias,t*_ = (−1.42, 0.075) for scenario (2) data points. In order to understand the source of these biases, we ran PAPI on the exact ancestry tracts output by the simulator for data from both scenarios. This analysis yields reduced bias measures of mean *d*_*bias,t*_ = (0.041, −0.701) for scenario (1) and *d*_*bias,t*_ = (−1.06, 0.394) for scenario (2) samples (Figure 5C,D), suggesting that some proportion of the downward bias can be explained by errors in local ancestry inference.

We contrasted these results, which are based on local ancestry estimates from HAPMIX, with those using tracts inferred by LAMP-LD (after filtering out tracts smaller than 1 cM; Methods). As shown in Supplementary Figure 5, the LAMP-LD-based estimates are biased upwards on average for both scenarios, with mean *d*_*bias,t*_ = (1.07, −0.089) for scenario (1) and *d*_*bias,t*_ = (−0.576, 1.66) for scenario (2) data points. The LAMP-LD-based estimates also show weaker correlations with the truth compared to the HAPMIX-based estimates with *R* = 0.59 for scenario (1) (considering only *t*_*B*_ values) and *R* = 0.69 for scenario (2) data points. Given these results, we conducted all further analyses using local ancestry tracts inferred by HAPMIX.

The above results consider the 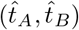 estimates in aggregate across simulated individuals, but PAPI is designed to be applied to individual samples as well. We plotted sample-specific point estimates of (*t*_*A*_, *t*_*B*_) (obtained using the MCMC inference mode) in the scenario (2) data along with their 90% Bayesian credible intervals (CIs) in Supplementary Figure 6. 90% CIs overlap true values in 86% of data points and have a mean range of 4.99. Notably, when the true *t*_*A*_ = *t*_*B*_ = 0, the 90% CIs of the model (in this case run with the error model option—see “Effectiveness of PAPI’s error model”) estimates have very large mean range of 16.5 and are strongly biased (*d*_*bias,t*_ = 7.06). This is due to the fact that when *t*_*A*_ = *t*_*B*_ = 0, both *p*_*A*_ and *p*_*B*_ must be either 0 or 1, and hence the inferred values in 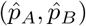 are each close to one of these extremes. Equation (2) implies that when this happens, either 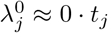 or 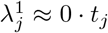, and therefore relatively small changes in *λ*_*j*_ can lead to large changes in *t*_*j*_; thus small erroneous tracts can readily bias the results. As noted below in the discussion of the error model, caution is advised in interpreting the *t*_*j*_ estimates when *p*_*j*_ *>* 0.95 or *p*_*j*_ *<* 0.05.

We further validated PAPI’s *t*_*j*_ estimates in real data by again analyzing the 10 ASW trio offspring. Here, the parents’ true time since admixture cannot be directly ascertained, so we first ran PAPI on each trio parent and computed the truth as an average of that parent’s time since admixture estimates, adding one for the meiosis to the trio offspring: 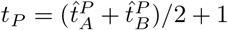 and 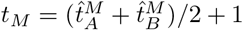, where for 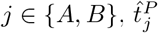 and 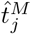 represent PAPI’s estimates on the trio father and trio mother, respectively. We then assigned the trio children’s 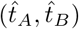 estimates to parent labels by minimizing the sum of squared differences (Methods). PAPI’s estimates in these data have a strong correlation with the truth of *R* = 0.77, in line with that observed in the simulated data (Figure 4B). These estimates and are somewhat biased with *d*_*bias,t*_ = (2.061, −0.332).

### Interpreting time since admixture estimates

As described in Methods, we simulated single haplotypes under a larger class of pedigree structures defined by four parameters—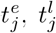 and *q*. We carried out these simulations with 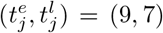 for each of *E*(*p*_*j*_) ∈ {0.25, 0.5}, where, when *E*(*p*_*j*_) = 0.5, *q* varied between [0, 1] in increments of 1*/*2^6^, and when *E*(*p*_*j*_) = 0.25, *q* varied between [0, 0.5] in the same increments (*q*’s maximum value is 0.5 for *E*(*p*_*j*_) = 0.25). We further simulated structures with 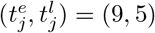 for *E*(*p*_*j*_) = 0.5, varying *q* between [0, 1] in increments of = 1*/*2^4^.

Given the exact local ancestry segments output by the simulation pipeline, we applied Equation (10) to estimate 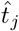 and then empirically examined the relationship between 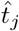 and the simulation parameters 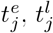, *E*(*p*_*j*_) and *q*.

Our results suggest that 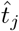 linearly increases from 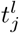 to 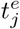 as *q* increases from 0 to its maximum value (Figure 6; Supplementary Figure 8), but notably, the 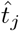 estimates are downwardly biased from the weighted average over all *N* simulated datapoints. In particular, when *E*[*p*_*j*_] = 0.5, this weighted average is 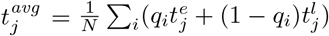, and when *E*[*p*_*j*_] = 0.25 we use 2*q*_*i*_ and (1 − 2*q*_*i*_) as the weights, with the factor of 2 accounting for *q*’s smaller range in this case. We measured the bias as 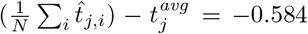 when *E*[*p*_*j*_] = 0.25 and −0.282 when *E*[*p*_*j*_] = 0.5. Similar biases arise in PAPI’s *t*_*j*_ estimates when using both simulated exact ancestry tracts and tracts inferred by HAPMIX (see previous subsection). While these results indicate that the estimated times since admixture may generally be a weighted average of the times in which individuals of a given ancestry entered an individual’s pedigree, a full understanding of the quantitative relationship requires more detailed analyses that are beyond the scope of this work. Instead we note simply that PAPI’s admixture time estimates potentially represent inputs from several generations and should not be interpreted as *the* generation in which admixture occurred.

**Figure 6:**
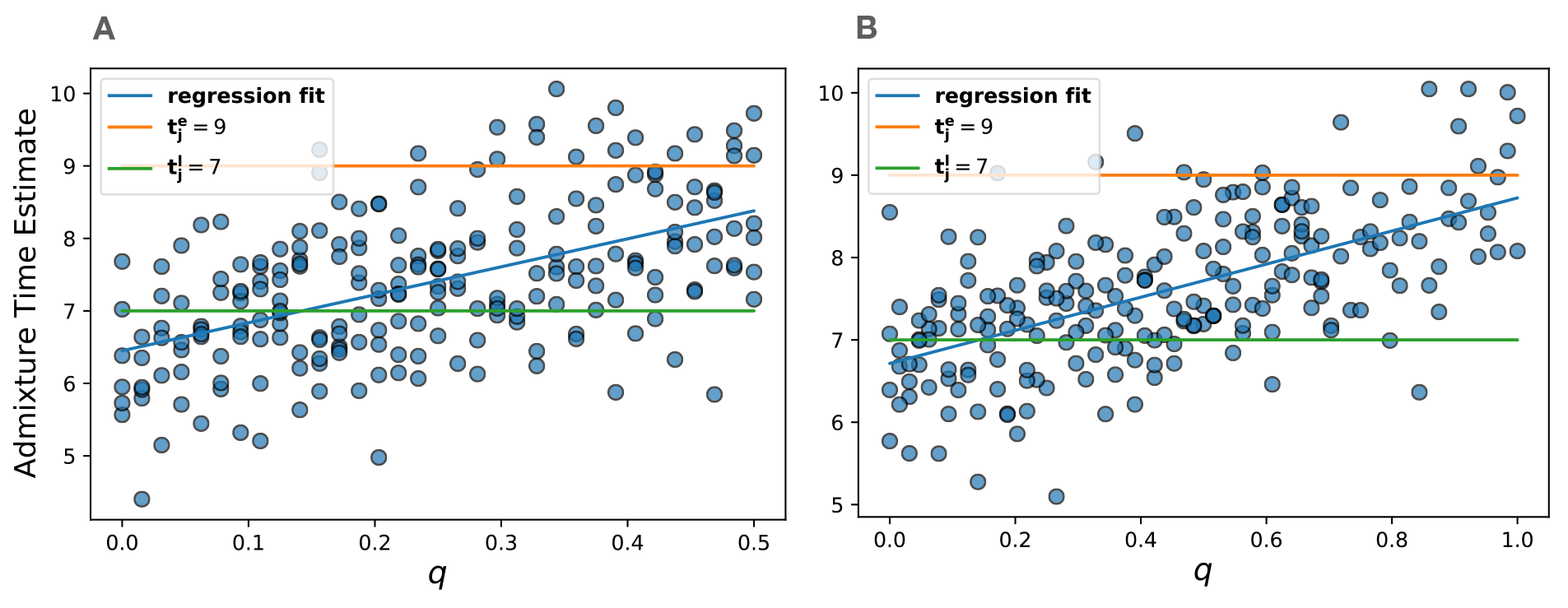
PAPI’s estimated time since admixture with two migrant pulses. Plots show estimated time since admixture versus *q* (the proportion of couples in the 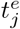 generation with unadmixed individuals of different ancestries) when 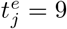 and 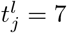 for (A) *E*[*p*_*j*_] = 0.25 and (B) *E*[*p*_*j*_] = 0.5.

### Effectiveness of PAPI’s error model

While generally reliable, local ancestry inference methods do produce erroneous tracts, which can lead to biased (*t*_*A*_, *t*_*B*_) estimates (see “Accuracy of time since admixture estimates”). These biases are most pronounced when the true *p*_*A*_ and *p*_*B*_ values are close to 0 or 1, such as in a subset of the scenario (3) simulated data (where *E*[(*p*_*A*_, *p*_*B*_)] ∈ {(0, 0), (1, 0), (1, 1)} and (*t*_*A*_, *t*_*B*_) = (0, 0)). In these cases, PAPI’s error model helps account for any erroneous tracts, leading to improved estimates of (*p*_*A*_, *p*_*B*_) (*d*_*abs,p*_ = 0.0078 with the error model as opposed to *d*_*abs,p*_ = 0.0148 without) and (*t*_*A*_, *t*_*B*_) (*d*_*bias,t*_ = 0.957 with the error model and *d*_*bias,t*_ = 9.934 without). Despite these improvements when the true *t*_*A*_ = *t*_*B*_ = 0, note that few small tracts that this model considers to be erroneous may in fact be real and reflect old admixture. As such, by default, PAPI runs without the error model, and we advise caution in interpreting its *t*_*j*_ estimates when *p*_*j*_ *>* 0.95 or *p*_*j*_ *<* 0.05. If a user is confident that the population under study is only recently admixed, using the error model can improve the results. In either case, it may be prudent to re-run PAPI in MCMC mode and examine the posterior distribution of (*t*_*A*_, *t*_*B*_) for samples with *p*_*j*_ *>* 0.95 or *p*_*j*_ *<* 0.05, or to avoid consideration of results from such parents.

### Ancestry proportions of parental couples are strongly correlated with each other in PAGE African Americans

We ran PAPI on detected African Americans from the BioMe Biobank subset of the PAGE study^27^, considering a total of 5,786 individuals that met our ancestry-based criteria for inclusion (Methods). Here, we initially examined the parental ancestry proportion estimates from PAPI’s binomial model run in MCMC inference mode—the recommended mode when only *p* estimates are required (see “Evaluating the performance of PAPI’s component models”). The average African ancestry proportion in the parents is *p* = 0.71, which is roughly consistent with previous estimates in African Americans^7,29,30^. The majority of parents have proportions between 0.8 and 1.0, corresponding to high African ancestry, but a heavy left tail in the distribution drives down the mean (Figure 7A). Figure 7B plots the estimated time since admixture—obtained by running PAPI with the default options—considering all parents that have *p*_*j*_ ∈ [0.05, 0.95] for *j* ∈ {*A, B*}. This distribution has an overall Gaussian-like shape, which may suggest random mating with respect to the time since admixture among the ancestors of these African Americans. The population mean of *t*_*j*_ estimates is 6.10, again roughly consistent with previous estimates in African Americans^7,29,30^. We infer a small set of individuals (0.04% of the dataset) with *t*_*j*_ *>* 15, but since the simulated data also contain some overestimated extreme values, we assumed that these estimates are inflated.

**Figure 7:**
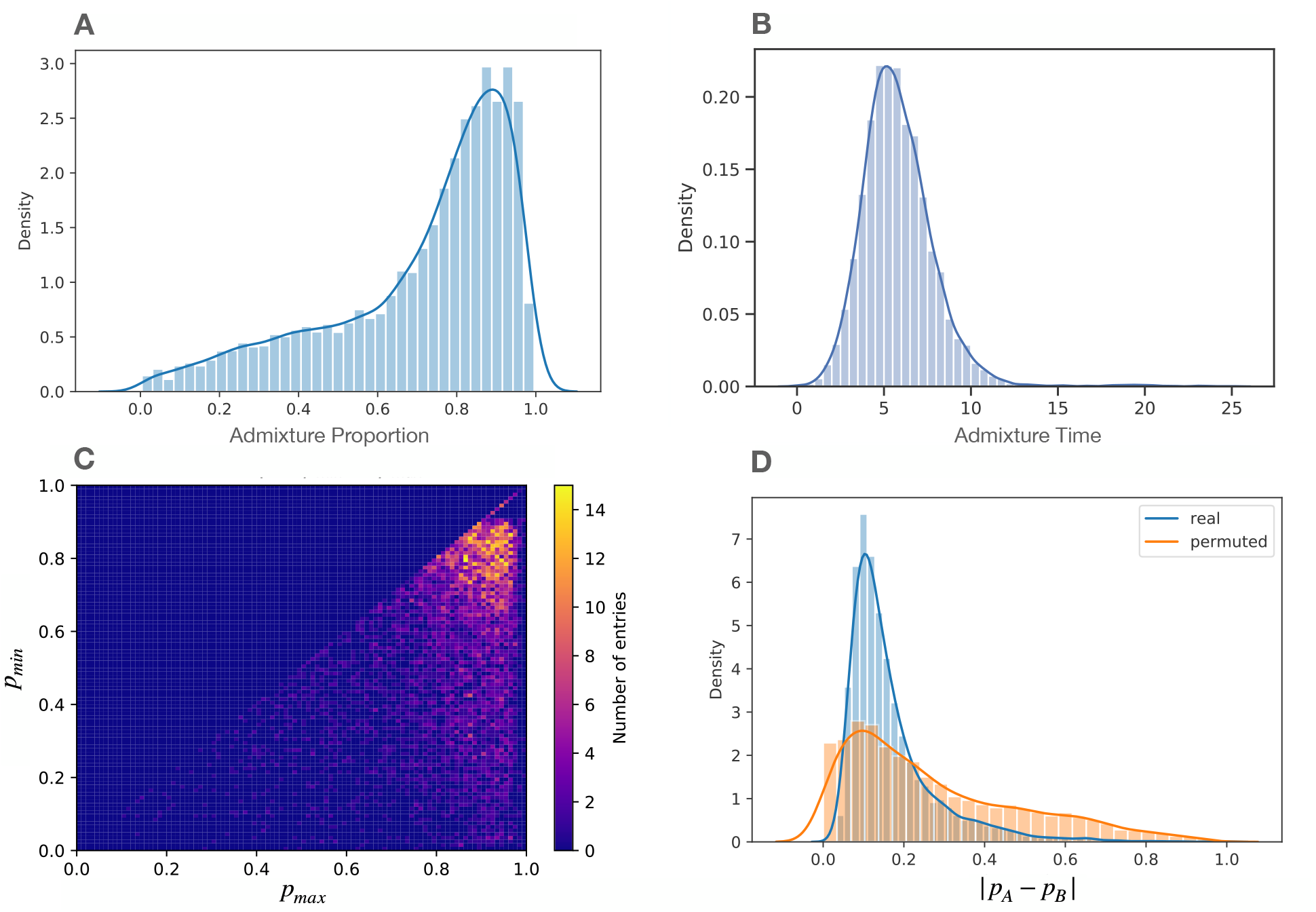
PAPI analyses from African Americans in the BioMe Biobank subset of PAGE. (A) Distribution of the parent ancestry estimates (both *p*_*A*_ and *p*_*B*_ plotted separately) in these African Americans. (B) Distribution of time since admixture estimates (*t*_*A*_ and *t*_*B*_ plotted separately). (C) Heatmap of the ancestry proportions of couples (*p*_*A*_, *p*_*B*_) ordered as (*p*_*max*_, *p*_*min*_). (D) Distribution of |*p*_*A*_−*p*_*B*_| estimates from the real couples along with a permuted distribution.

To leverage PAPI’s ability to estimate parental admixture proportions, we analyzed these estimates within all parent pairs—all couples—from these African Americans. Figure 7C depicts a heatmap of (*p*_*max*_, *p*_*min*_) = (max(*p*_*A*_, *p*_*B*_), min(*p*_*A*_, *p*_*B*_)) (since the *A* and *B* labels are arbitrary) where one salient feature is a strong cluster of points centered roughly at (0.95, 0.85) that spreads diffusely outward. This cluster is consistent with a pattern of assortative mating between individuals with high African ancestry, and includes a meaningful fraction of the couples (25.2% of individuals in the dataset have *p*_*max*_ ≥ 0.90 and *p*_*min*_ ≥ 0.80). Furthermore, the Pearson correlation coefficient between *p*_*max*_ and *p*_*min*_ is *R* = 0.871, also indicating assortative mating by ancestry.

To more precisely characterize the strength of the signal for non-random mating in these couples, we examined the per-individual distribution of |*p*_*A*_ − *p*_*B*_| (Figure 7D). This density is roughly Gaussian with a heavy right tail, and has a mean of 0.169, largely driven by the high density of couples noted above. This real data distribution is visibly distinct from what would arise under random mating, as randomly shuffling the members of the couples creates a much more dispersed, thick-tailed density (Figure 7D). Indeed, the mean |*p*_*A*_ − *p*_*B*_| across 10^6^ permutations is 0.272, and no permutation mean reached as small a value as in the real data (minimum permutation mean 0.264; *P <* 10^−6^). Taken together, these observations provide strong evidence for assortative mating by ancestry proportion in the parents of these African American individuals.

## Discussion

In this paper, we described and evaluated PAPI, a novel approach to inferring the admixture proportions and times since admixture for each parent of an extant individual that combines two models. The first is a binomial model that takes a global view of the ancestry of an individual, considering the overall proportions of homozygous and heterozygous ancestry in their genome. As noted earlier, this model is equivalent to that described in a recent manuscript^28^. PAPI employs a Bayesian framework and combines this binomial model with a second HMM that leverages the information present in the length distribution of local ancestry tracts. This HMM incorporates a population genetic model^9^—but uses parameters specific to the individual’s pedigree—to capture the hidden recombinations within these tracts (i.e., those that do not switch the ancestry of the haplotype).

To characterize PAPI’s accuracy, we simulated African Americans under various pedigree topologies and distributions of parent ancestry proportions and also used the real HapMap ASW trios. Our results show that PAPI is more accurate than ANCESTOR at inferring parent ancestry proportions in many settings (Figure 4) and can detect when *t*_*A*_ ≠ *t*_*B*_ (Figure 5). In simulations that rely on inferred local ancestry tracts, PAPI is able to recover *t*_*j*_ estimates highly correlated with the truth, albeit with a downward bias when using HAPMIX local ancestry tracts and an upward bias when using LAMP-LD (Figure 5).

PAPI’s approach models historical crossovers using a Markovian approximation to the more elaborate dynamics of crossover transmissions in a pedigree. This is similar to ANCESTOR’s HMM, although ANCESTOR also supports non-exponential tract length distributions using a stochastic expectation maximization technique for inference^37^. In contrast, PedMix^15^ parameterizes the full pedigree up to the grandparental generation using an ancestral configuration vector (which captures grandparental ancestry states, phase error, and all transmitted recombinations) while treating older recombinations as Markovian. Despite only approximating pedigree-based crossovers, and through its two component models, PAPI produces highly accurate estimates of parental ancestry proportions without relying on phased input. Note that neither PedMix nor ANCESTOR estimate parental admixture times (ANCESTOR instead provides estimates of average ancestry tract lengths for each parent).

PAPI’s ancestry estimates in the PAGE dataset add to a growing body of evidence for assortative mating by ancestry proportion in African Americans^22^. Others have noted these signals in Latinos^21^ and found that they frequently cannot be explained by socioeconomic factors alone, and are important to take into account when designing genetic studies. For example, using Wright-Fisher simulations that allow for ancestry-based assortative mating, Zaitlen et al.^22^ showed that assuming random-mating leads to significant underestimates of population-scale admixture times.

Overall, PAPI’s applications include both the study of the population genetics of mating in African Americans, and the potential to provide individual-level information to study subjects. Given this ability to infer parental ancestry proportions for individual samples, we expect PAPI to be of interest to direct-to-consumer genetic testing companies and to consumers themselves who wish to better understand their genetic heritage.

## Acknowledgements

Funding for this work was provided by NIH grant R35 GM133805. Computing was performed on a cluster administered by the Biotechnology Resource Center at Cornell University.

## Data and code availability

The PAPI code is available at https://github.com/williamslab/papi. Genotype data for BioMe Biobank subset of the PAGE II dataset are available (dbGaP:phs000925.v1.p1), and the phased 1000 Genomes data used for local ancestry inference is publicly available at https://www.internationalgenome.org/data-portal/sample. The simulated genotype data supporting the current study have not been deposited in a public repository but can be reproduced using Ped-sim, admix-simu and the publicly available HapMap CEU and YRI data (https://ftp.ncbi.nlm.nih.gov/hapmap/).

