## Supplementary Data for "Simultaneous inference of parental admixture proportions and admixture times from unphased local ancestry calls"

| Model Setting / Inference Mode | Binomial | Binomial w/Error Model | HMM | HMM w/Error Model | Full | Full w/Error Model |
|---|---|---|---|---|---|---|
| **GD** | 7.3s | – | 21.7s | 23.6s | 23.9s | 27.0s |
| **MCMC** | 5m 23s | – | 8h 36m 49s | 8hr 58m 31s | 8hr 5m 58s | 8hr 55m 6s |

Table S1: **PAPI runtimes for each model and inference mode.** Times are the per individual average from simulated samples with $t_A = t_B = 5$, including 22 individuals for each of $E[(p_A, p_B)] \in \{(0.5, 0.5), (0.25, 0.5), (0.4, 0.6), (0.25, 0.75)\}$, for a total of 88 individuals per run. Table gives wall clock times from compute nodes with Intel Xeon E5 4620 processors utilizing 4 GB of RAM.

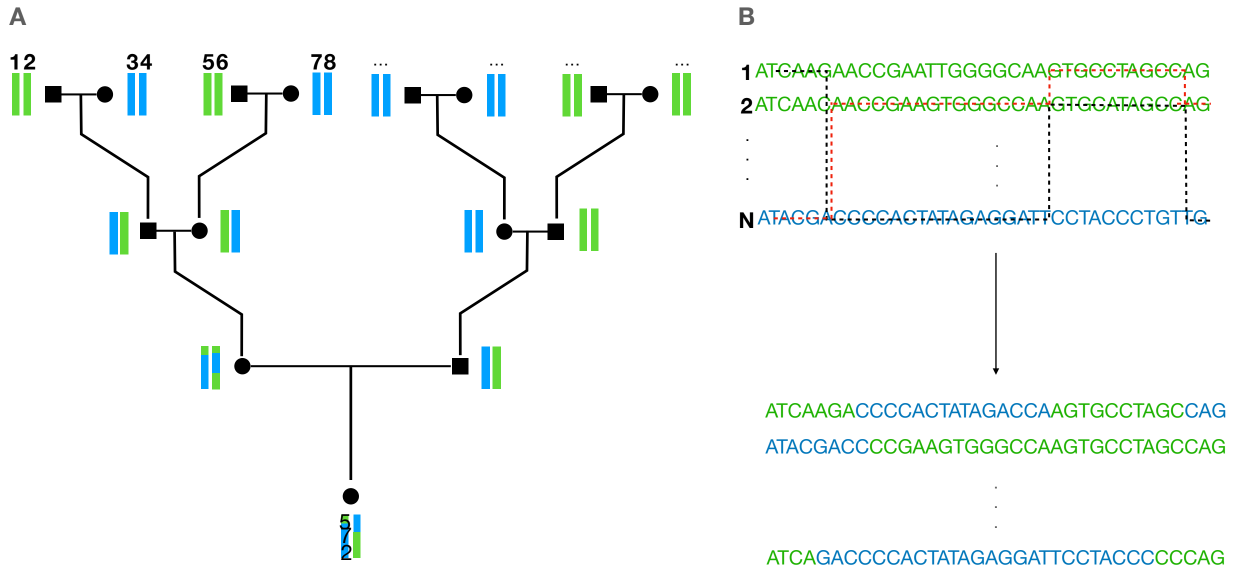


Figure S1: **Overview of simulation procedure.** (A) Given a pedigree topology (specified by $(t_A, t_B)$), Ped-sim generates haplotype records for the simulated samples containing ancestral recombination break points and a numerical founder haplotype id for each segment. Subsequently, we assign these haplotype ids to population labels according to the desired values of $E[(p_A, p_B)]$ and $(t_A, t_B)$. In the example depicted here, $(t_A, t_B) = (2, 1)$ and $E[(p_A, p_B)] = (0.5, 0.5)$, where parent $A$ is on the left and $B$ is on the right. (B) Given the adapted Ped-sim output containing population labels, admix-simu generates $N$ offspring genotypes by sampling non-overlapping paths (black and red dashed lines representing the sampling paths of two haplotypes) from $N$ founder YRI and CEU genotypes.

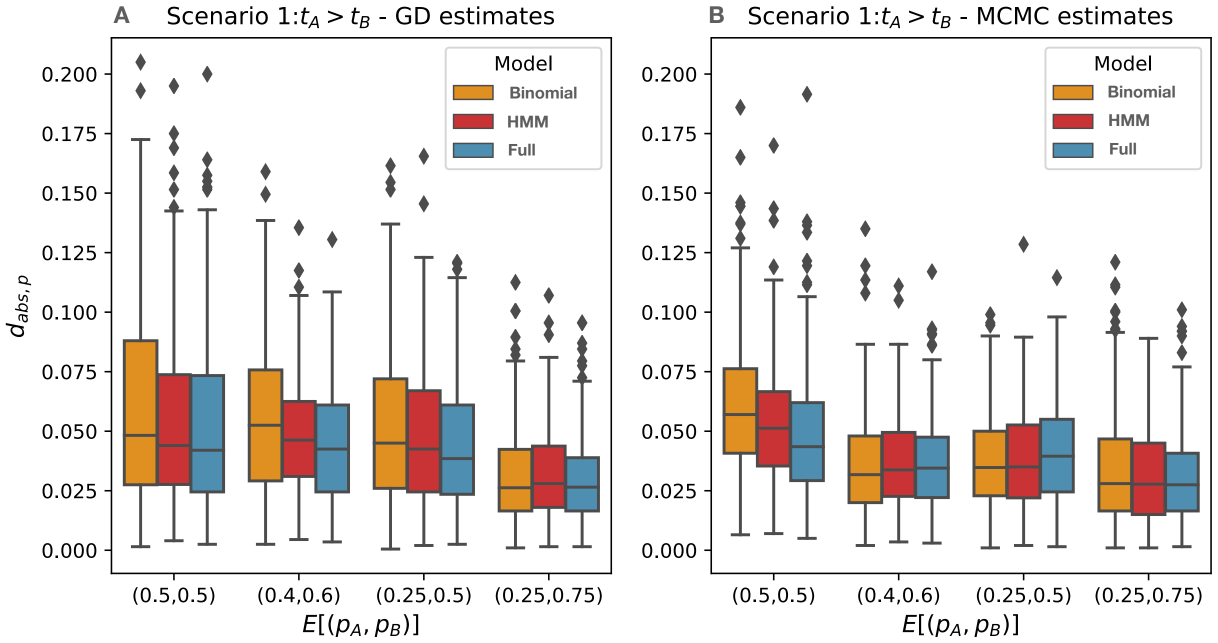


Figure S2: **Deviance statistics for inferred parent ancestry proportions** $(p_A, p_B)$ **for each of PAPI's component models.** Average absolute deviances $(d_{abs})$ on all scenario (1) data under the (A) GD and (B) MCMC inference modes.

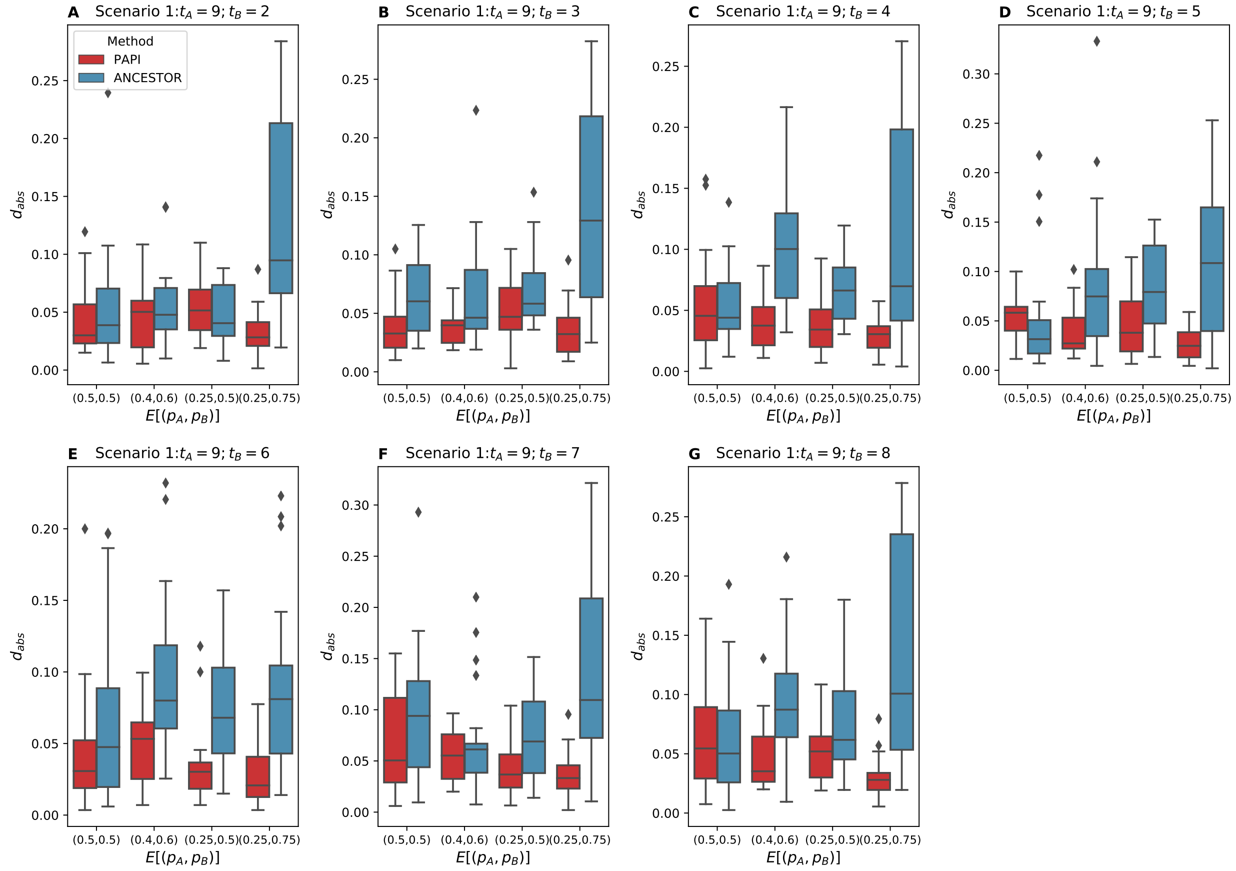


Figure S3: **Deviance statistics for inferred parent ancestry proportions** $(p_A, p_B)$ **from ANCESTOR and PAPI for individual scenario (1) data points.** (A-G) Average absolute deviances ($d_{abs}$) for each value of $(t_A, t_B)$ as indicated in the panel title.

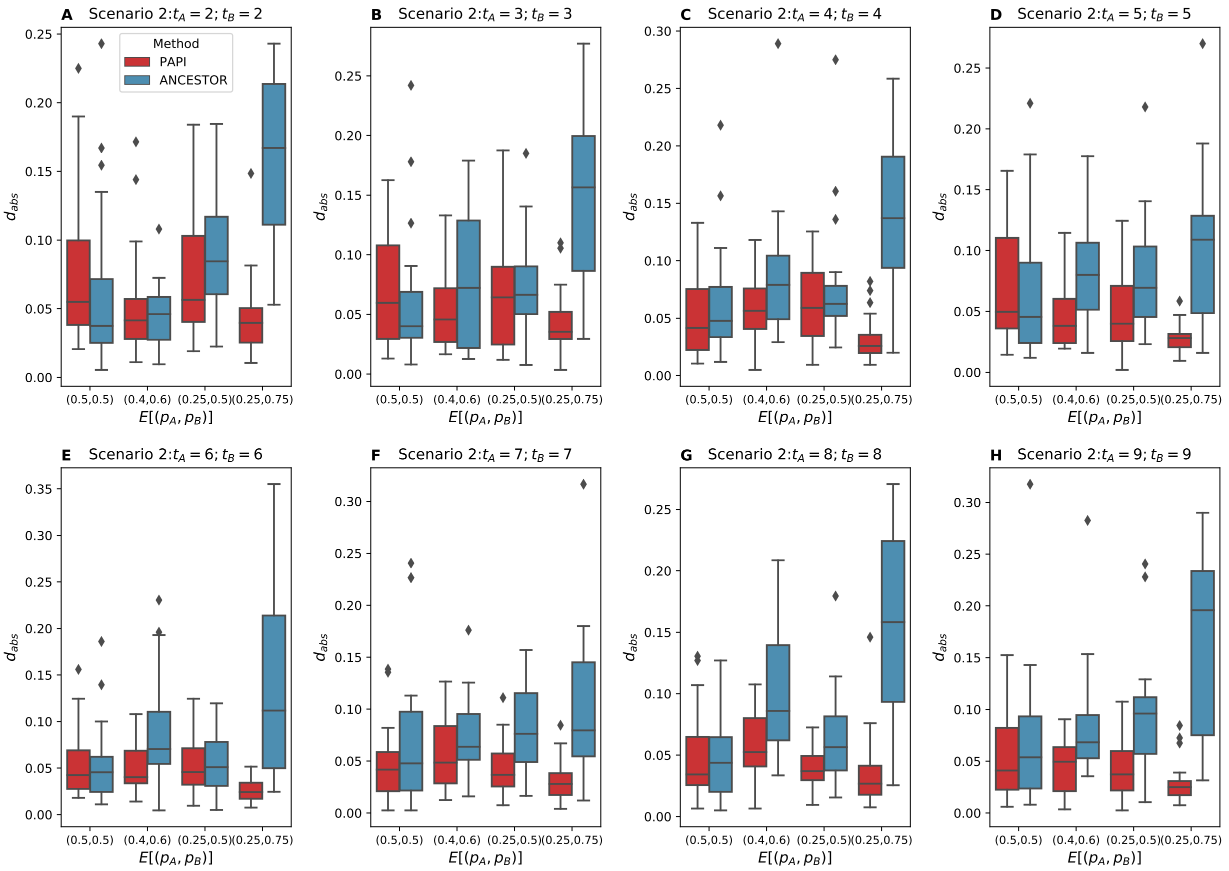


Figure S4: **Deviance statistics for inferred parent ancestry proportions** $(p_A, p_B)$ **from ANCESTOR and PAPI for individual scenario (2) data points.** (A-H) Average absolute deviances ($d_{abs}$) for each value of $(t_A, t_B)$ as indicated in the panel title.

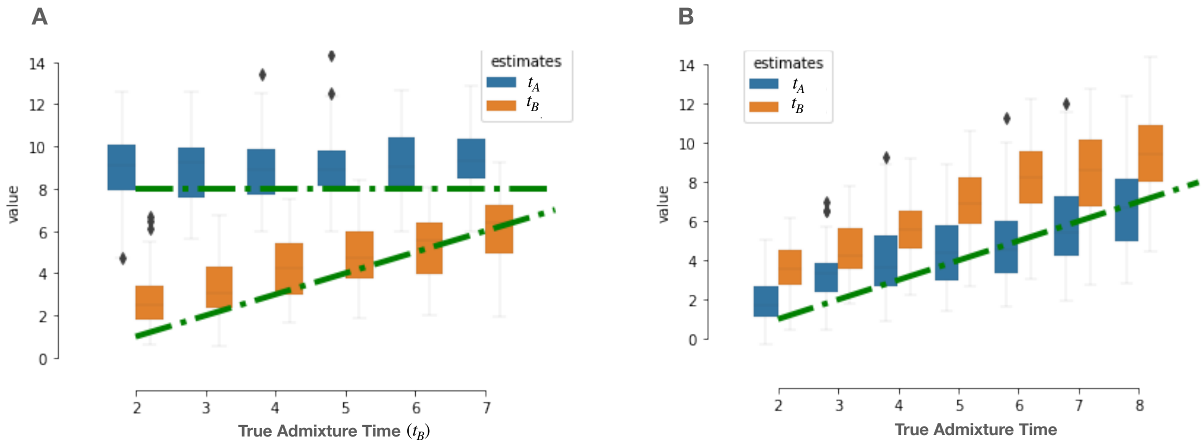


Figure S5: **PAPI's estimated time since admixture in simulated individuals using local ancestry tracts inferred by LAMP-LD** Box plots of estimated times for (A) scenario (1) samples (where $t_A > t_B$) and (B) scenario (2) data (where $t_A = t_B$).

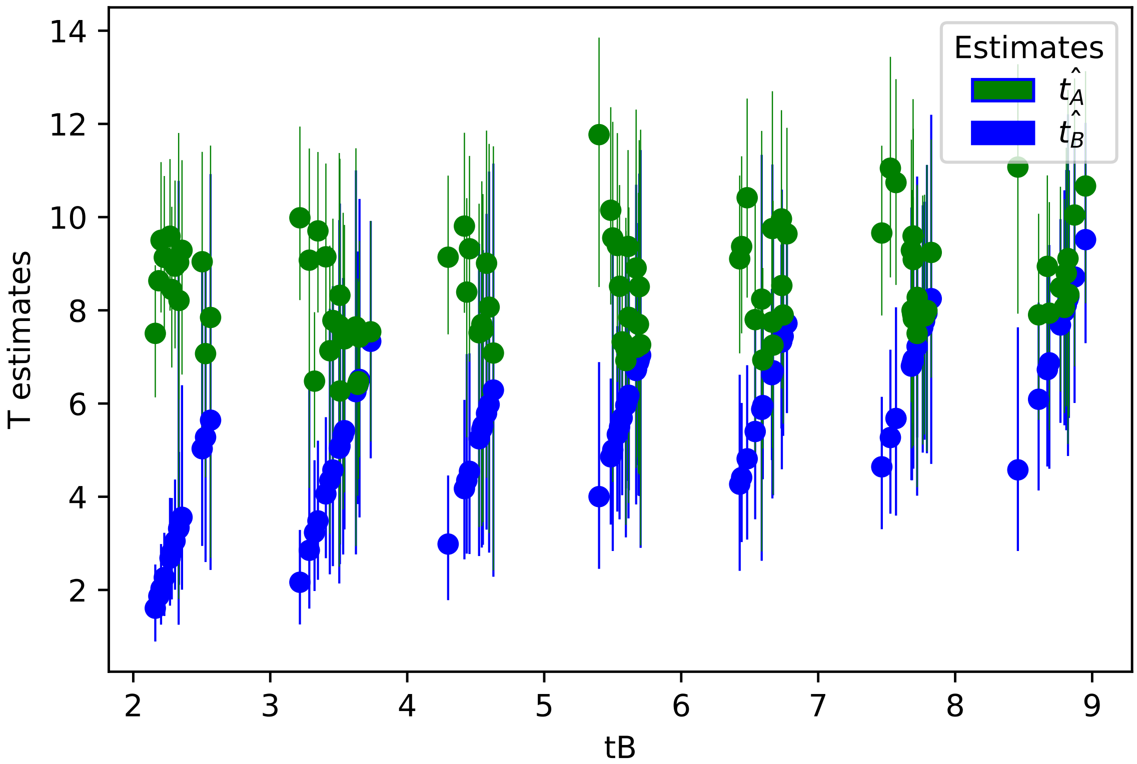


Figure S6: **Admixture time estimates with credible intervals for scenario (1) data points.** Plot shows point estimates along with their 90% credible intervals from PAPI's MCMC inference mode run under the full model. The x-axis coordinate for each point is shifted slightly from the truth to aid visualization: the estimate pairs $(\hat{t}_A, \hat{t}_B)$ are sorted by the value of $\hat{t}_B$.

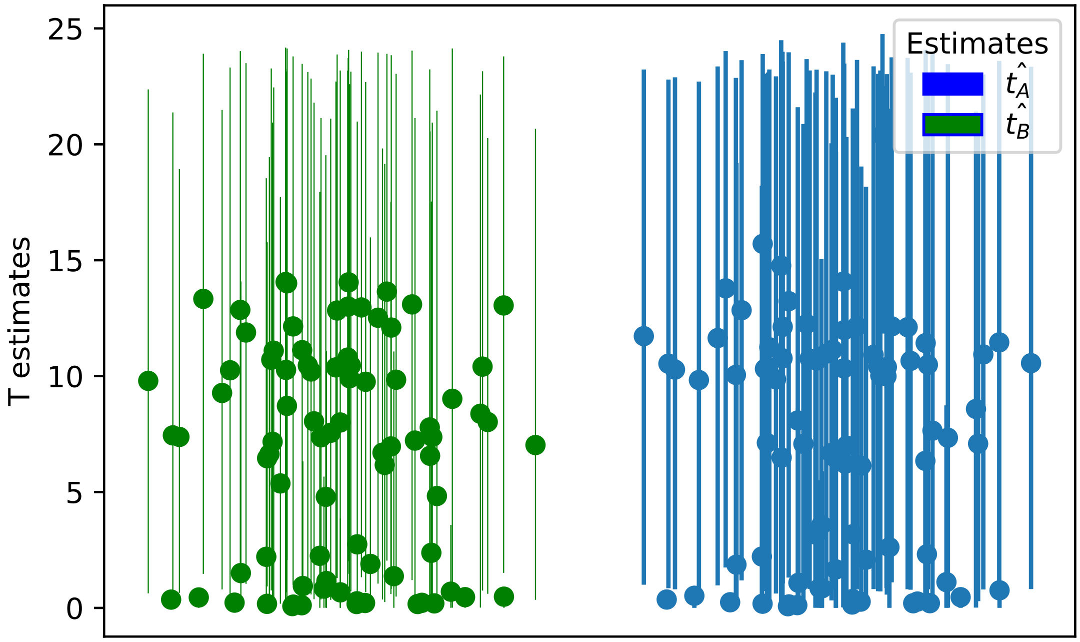


Figure S7: **Admixture time estimates with credible intervals for the subset of scenario (3) data points with $t_A = t_B = 0$.** Plot shows point estimates along with their 90% credible intervals from PAPI's MCMC inference mode run under the full model. Many estimates are biased and have very large 90% credible intervals because of a small number of erroneous tracts.

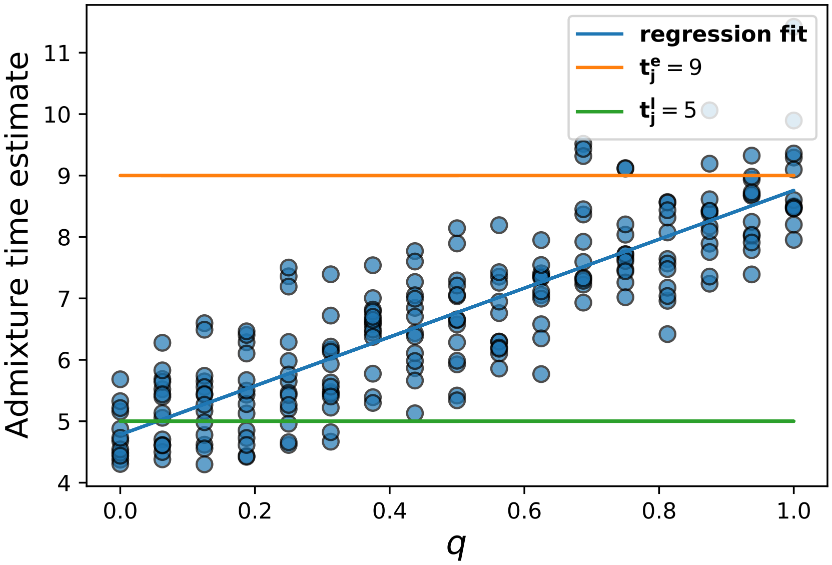


Figure S8: **PAPI's estimated time since admixture with two migrant pulses with $t_j^e = 9$ and $t_j^l = 5$.** Plot of estimated time since admixture versus q (the proportion of unadmixed couples in the $t_j^e$ generation of different ancestries) when $E[p_j] = 0.5$.
